# Direct cell-to-cell transmission of retrotransposons

**DOI:** 10.1101/2025.03.14.642691

**Authors:** Maya Voichek, Andreas Bernhard, Maria Novatchkova, Dominik Handler, Paul Möseneder, Baptiste Rafanel, Peter Duchek, Kirsten-André Senti, Julius Brennecke

## Abstract

Transposable elements are abundant in host genomes but are generally considered to be confined to the cell in which they are expressed, with the notable exception of endogenous retroviruses. Here, we identify a group of LTR retrotransposons that infect the germline from somatic cells within the *Drosophila* ovary, despite lacking the fusogenic Envelope protein typically required for retroviral entry. Instead, these elements encode a short transmembrane protein, sORF2, with structural features reminiscent of viral cell-cell fusogens. Through genetics, imaging, and electron microscopy, we show that sORF2 localizes to invasive somatic protrusions, enabling the direct transfer of retrotransposon capsids into the oocyte. Remarkably, sORF2-like proteins are widespread among insect retrotransposons and also occur in piscine nackednaviruses and avian picornaviruses. These findings reveal a noncanonical, Envelope-independent transmission mechanism shared by retrotransposons and non-enveloped viruses, offering important insights into host-pathogen evolution and soma-germline interactions.

## INTRODUCTION

Viruses are “master hackers” of cell biology and understanding the molecular determinants that underlie their life cycles, particularly cell-to-cell transmission, has important medical and scientific implications. While much of what we know about viral entry and exit mechanisms comes from studying modern human pathogens, these represent only a small fraction of the viral realm. Throughout evolution, most virus-host battles have been won or lost without leaving a trace, but vertebrate endogenous retroviruses (ERVs) are a fascinating exception. These genomic fossils are thought to have arisen from rare germline infections of exogenous retroviruses that led to the integration of the viral genome into the host genome and are thus often grouped together with transposable elements (TEs), specifically long terminal repeats (LTR) retrotransposons (Johnson, 2019). The repertoire of ERVs present in a host genome hence captures a “snapshot” of past infections, providing a unique opportunity to study the molecular basis of viral infectivity. However, ERVs degenerate over time by accumulating mutations due to powerful repression mechanisms against selfish genetic elements in the host germline (Levin & Moran, 2011). Together with the high copy number of ERVs in vertebrate genomes, studying their biology and in particular their infectivity poses significant challenges.

The fruit fly *Drosophila melanogaster* offers an alternative promising model for studying endogenous retroviruses in the context of germline biology. Its genome harbors a wealth of relatively young, diverse and, above all, active ERVs (Kaminker et al., 2002; Kapitonov & Jurka, 2003). While insect ERVs (family *Metaviridae*) are phylogenetically distinct from vertebrate ERVs (family *Retroviridae*), some share key structural and functional features, including the canonical retroviral genome organization with *gag*, *pol* and *env* genes flanked by two LTRs (fig. S1A) (Llorens et al., 2020; Malik et al., 2000). Under permissive conditions, enveloped insect-ERVs such as *Gypsy* or *ZAM* are expressed in somatic cells of the ovary and infect the neighboring oocyte as retroviral particles (A. Kim et al., 1994; Leblanc et al., 2000; Song et al., 1994; Yoth et al., 2023). Similar to vertebrate retroviruses, this cell-to-cell infectivity is thought to require an intact *envelope* (*env*) gene, which encodes a fusogenic protein mediating the virus-cell membranes fusion (Misseri et al., 2004; Pélisson et al., 1994). In contrast, many other LTR retrotransposons in the *Drosophila* genome lack a functional *env* gene and are expressed in the germline lineage like other TEs (Bourque et al., 2018). Both insect ERVs and LTR retrotransposons are suppressed by the host defense system, the PIWI/piRNA pathway, which is active in the gonad (Malone & Hannon, 2009; Senti et al., 2023). Upon genetic disruption of the piRNA pathway, active insect ERVs and LTR retrotransposons are expressed, creating an opportunity to investigate their native biology. Utilizing the rich toolkit of *Drosophila* genetics, we set out to pursue the possibility of novel features that confer infectivity to LTR retrotransposons.

In this study, we uncover an *env*-independent transmission mechanism that enables a group of LTR retrotransposons, the MDG1 elements, to directly enter the well-protected oocyte from neighboring somatic cells. This discovery highlights an evolutionary innovation by which an ancestral LTR retrotransposon has gained viral-like cell-to-cell transmission capability, further blurring the conventional boundaries between transposons and viruses (Widen et al., 2023). Remarkably, we find that similar *env*-independent infectivity strategies are prevalent among non-enveloped viruses infecting a broad spectrum of hosts.

## RESULTS

### LTR retrotransposons of the MDG1 group behave as retroviruses

LTR retrotransposons are generally confined to the cells in which they are expressed, and in animals their evolutionary survival strategy therefore relies on expression and generation of new genome insertions in the germline. A notable exception is found in the *Gypsy* group of *Metaviridae*, in which some members encode an *env* gene (Fig. 1A). Previous work has demonstrated a strong correlation between a functional *env* gene and exclusive expression in somatic gonadal cells, enabling these elements to infect the germline. In contrast, *Gypsy* LTR retrotransposons lacking an *env* gene are expressed specifically in the germline (Senti et al., 2023).

**Figure 1:**
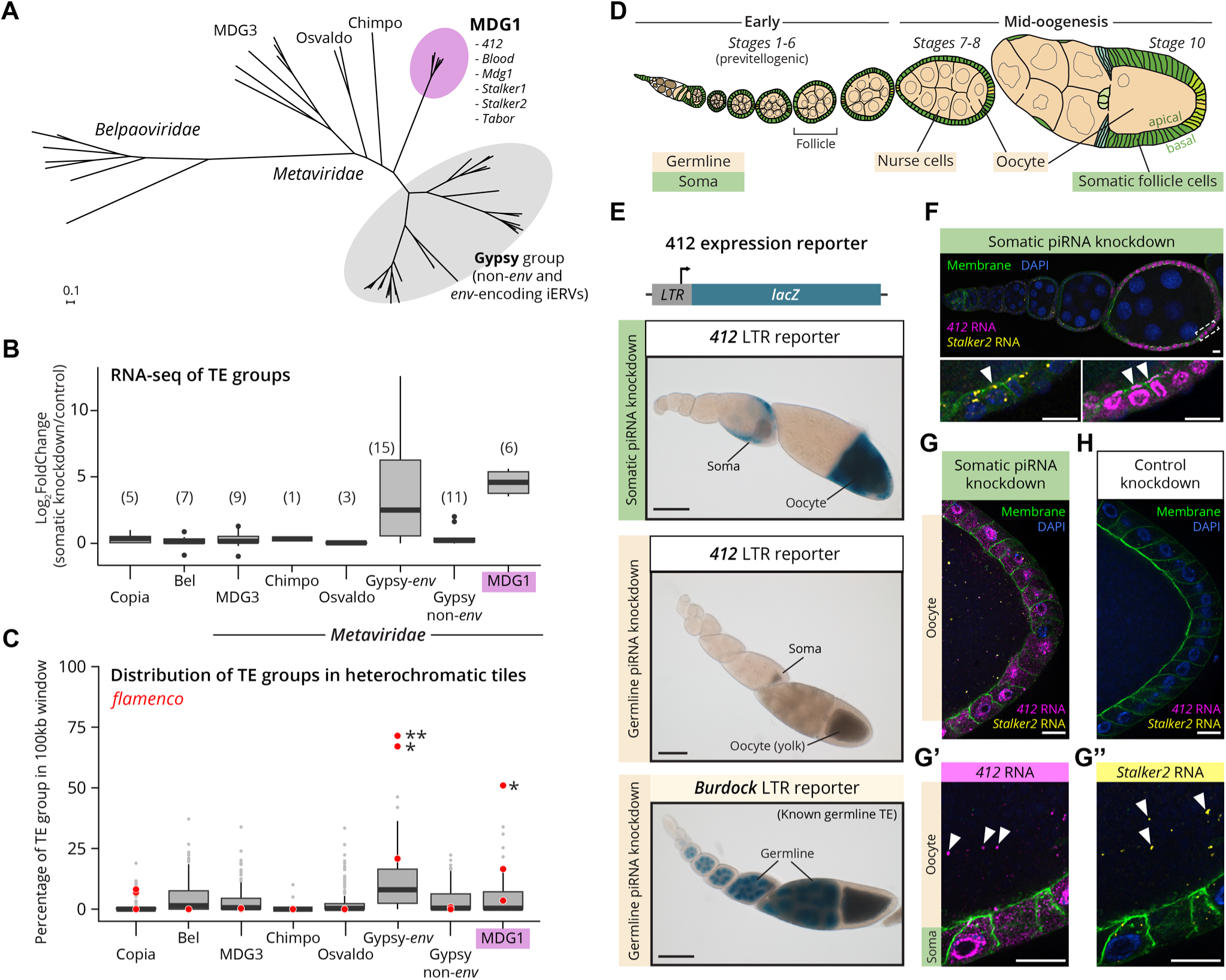
Somatically expressed *412* and *Stalker2* RNA transmits into the oocyte. **(A)** Phylogenetic tree of *Metaviridae* in *D. melanogaster* based on a multiple sequence alignment of the consensus *pol* region, rooted with *Belpaoviridae* as an outgroup. The Gypsy group (grey circle) includes both *env*-encoding insect ERVs and LTR retrotransposons lacking an *env* gene. The MDG1 group is highlighted in magenta. Scale bar: substitutions per site. **(B)** Expression fold change of transposable element (TE) groups measured by RNA-seq of ovaries upon somatic piRNA pathway knockdown (*Tj>Gal4, vret-RNAi*) compared to control (*Tj>Gal4, arr2-RNAi*) (n=3). The number of TEs in each group is indicated in brackets. **(C)** TE group distribution within TE-enriched heterochromatic tiles (100-kb genomic regions with >50% transposon content). Red datapoints represent *flamenco* tiles. Asterisks indicate significant deviation from the background distribution (MDG1 *p*-value = 0.0145). For (B-C) Gypsy-*env* refers to *env*-encoding insect ERVs, and Gypsy non-*env* refers to LTR retrotransposons without an *env* gene. Boxplots in (B-C) present the median (center line), IQR (box bounds), min and max within 1.5 IQR (whiskers), and outliers (points beyond whiskers). **(D)** Schematic of a *Drosophila* ovariole, showing sequential progression of follicles through the developmental stages of oogenesis in the adult ovary. Somatic follicle cells (green) are separate from the germline (beige), which is comprised of the oocyte and nurse cells. **(E)** Transcriptional *LTR-lacZ* reporters for *412* under different genetic conditions, (top) somatic piRNA pathway knockdown (*Tj>Gal4, vret-RNAi*); (middle) germline piRNA pathway knockdown (*MTD>Gal4, aub-RNAi, ago3-RNAi*); (bottom) *LTR-lacZ* reporter for germline-expressed *Burdock* TE under germline piRNA pathway knockdown (*MTD>Gal4, aub-RNAi, ago3-RNAi*). Scale bar: 100 µm. **(F-H)** Dual smFISH staining of early (F) and mid-oogenesis follicles (G-H) for LTR retrotransposons *412* (magenta) and *Stalker2* (yellow) RNA. Panels **(F)** and **(G, G’, G’’)** show somatic piRNA pathway knockdown (*Tj>Gal4, vret-RNAi*) while **(H)** shows the control knockdown for an unrelated gene (*Tj>Gal4, arr2-RNAi*). Panel (F) bottom shows zoom-in of boxed region in top image, at a different Z-section. White arrowheads denote RNA signals of LTR retrotransposons in the apical membrane of somatic cells in (F) and in the oocyte in (G’, G’’). Somatic membranes are marked by myristoylated GFP. Scale bars (F-H): 10 µm.

To identify additional LTR retrotransposons with potential soma-to-germline infectivity, we systematically analyzed their repertoire in *D. melanogaster* using two complementary approaches. First, we inactivated the piRNA pathway specifically in somatic ovarian cells using transgenic RNAi (*vreteno*-RNAi) (Handler et al., 2011), and determined LTR retrotransposon expression using RNA-seq of ovaries. As expected, *Gypsy* group elements with functional *env* were derepressed compared to control, while those lacking *env* were not (Fig. 1B) (Senti et al., 2023). However, all six elements from an additional *Metaviridae* group, the MDG1 clade, were also strongly derepressed by 24-fold despite lacking an *env* gene (Kapitonov & Jurka, 2003). Second, we analyzed the silencing spectrum of the piRNA pathway in the ovarian soma by examining the somatic master piRNA source locus, *flamenco* (Brennecke et al., 2007; Pélisson et al., 1994). Specifically, we asked whether insertions of any LTR retrotransposon group are enriched in *flamenco*. This analysis revealed that, in addition to *env*-encoding *Gypsy* group elements, one additional group is significantly enriched in *flamenco,* the abovementioned MDG1 clade (Fig. 1C). Taken together, LTR retrotransposons of the MDG1 group are expressed in somatic follicle cells under permissive conditions, and the host has evolved somatic piRNA defense against them.

To confirm that the somatic expression of MDG1 LTR retrotransposons is an intrinsic trait and not due to insertions affected by somatically expressed genes, we selected the most highly expressed MDG1 retrotransposon in piRNA pathway knockdown conditions, called *412*, for transcriptional reporter analysis. To this end, we cloned the *lacZ* coding sequence downstream of the LTR of *412*, which harbors the promoter and *cis*-regulatory elements required for expression (Johnson, 2019; Yuki et al., 1986). As expected, the reporter was not expressed in control ovaries, consistent with abundant *412* antisense piRNAs (fig. S1B). However, in ovaries with somatic piRNA pathway knockdown, we observed strong X-Gal staining exclusively in somatic follicle cells (Fig. 1D-E). In contrast, the reporter was not active in ovaries with defective germline piRNA pathway, as opposed to a reporter for a known germline-expressed LTR retrotransposon, *Burdock*, used as a positive control (Fig. 1E bottom) (Senti et al., 2023).

Given these observations, we hypothesized that somatically expressed MDG1 group retrotransposons evolved means to invade the germline, despite them lacking an *env* gene or the sequence space for an unknown open reading frame downstream of *pol* (fig. S1A). To test this, we used single molecule Fluorescence In Situ Hybridization (smFISH) to track the RNA genomes of MDG1 group retrotransposons in ovaries with a somatic piRNA pathway knockdown. During early oogenesis, abundant *412* RNA molecules were found exclusively in nuclei and cytoplasm of somatic follicle cells (Fig. 1F, fig. S1C). However, from stage 6-7 onward, *412* transcripts accumulated at the apical membrane of somatic follicle cells, close to the oocyte, and were also detected inside the oocyte in discrete foci despite an intact germline piRNA pathway (Fig. 1G). The presence of *412* transcripts in the oocyte was surprising, given the strict separation between germline and soma compartments with no cytoplasmic continuity in *Drosophila*. In control ovaries with intact piRNA/PIWI pathway, *412* RNAs were not detectable in somatic cells nor in the oocyte (Fig. 1H). Similar results were observed for the related elements *Stalker2* (Fig. 1F-H) and *Mdg1* (fig. S1D-F), suggesting that MDG1 group retrotransposons are capable of efficient soma-to-germline transmission.

Collectively, our findings indicate that *env*-less LTR retrotransposons of the MDG1 group have adapted viral traits, including somatic expression and a cell-cell transmission strategy, to target the oocyte genome from adjacent somatic cells, thus mimicking infectious retroviruses.

### The LTR retrotransposons 412 and Stalker2 exhibit autonomous infectivity

Retroviruses infect cells through their Envelope protein, a fusogenic transmembrane glycoprotein that binds to cell surface receptors and facilitates entry via viral-cell membrane fusion. We reasoned that the cell-to-cell transmission observed for MDG1 group retrotransposons might depend on complementation or pseudotyping by Envelope proteins of *Gypsy* group retroviruses, which are also expressed in piRNA deficient follicle cells (Barckmann et al., 2018; Teysset et al., 1998). To test this, we generated flies that are permissive exclusively for MDG1 group retrotransposons. In *D. melanogaster*, almost all transposon-silencing piRNAs in somatic cells originate from the master control locus *flamenco* (Brennecke et al., 2007; Malone et al., 2009). In the reference genome strain, *iso-1*, nested antisense insertions of the MDG1 group elements *412* and *Stalker2* are located ∼8kb downstream of the *flamenco* transcriptional start site, flanked by genome-unique sequence stretches (Zanni et al., 2013). Using CRISPR genome editing of the *iso-1* X-chromosome, we inserted two FRT sites flanking the *412*-*Stalker2* locus and excised the intervening 15.5kb sequence with the FLP recombinase (Fig. 2A). To validate the resulting *flamenco*^Δ*412*-*St2*^ allele, we sequenced ovarian piRNAs from two independently generated homozygous mutant lines. The targeted *flamenco* deletion did not change the piRNA pools targeting any transposon, except for *412* and *Stalker2*, whose antisense piRNA levels were reduced ∼15 and ∼30-fold, respectively (Fig. 2B; fig. S2A). At the same time, differential gene expression analysis based on poly(A)-enriched RNA-seq confirmed that *412* and *Stalker2* were the only de-repressed transposons with significantly increased sense transcript levels (∼50-fold; Fig. 2C; fig. S2B-C). The *flam^Δ412-St2^* allele is therefore an ideal tool to study MDG1 group retrotransposons without confounding impacts from *env*-encoding *Gypsy* group retroviruses.

**Figure 2:**
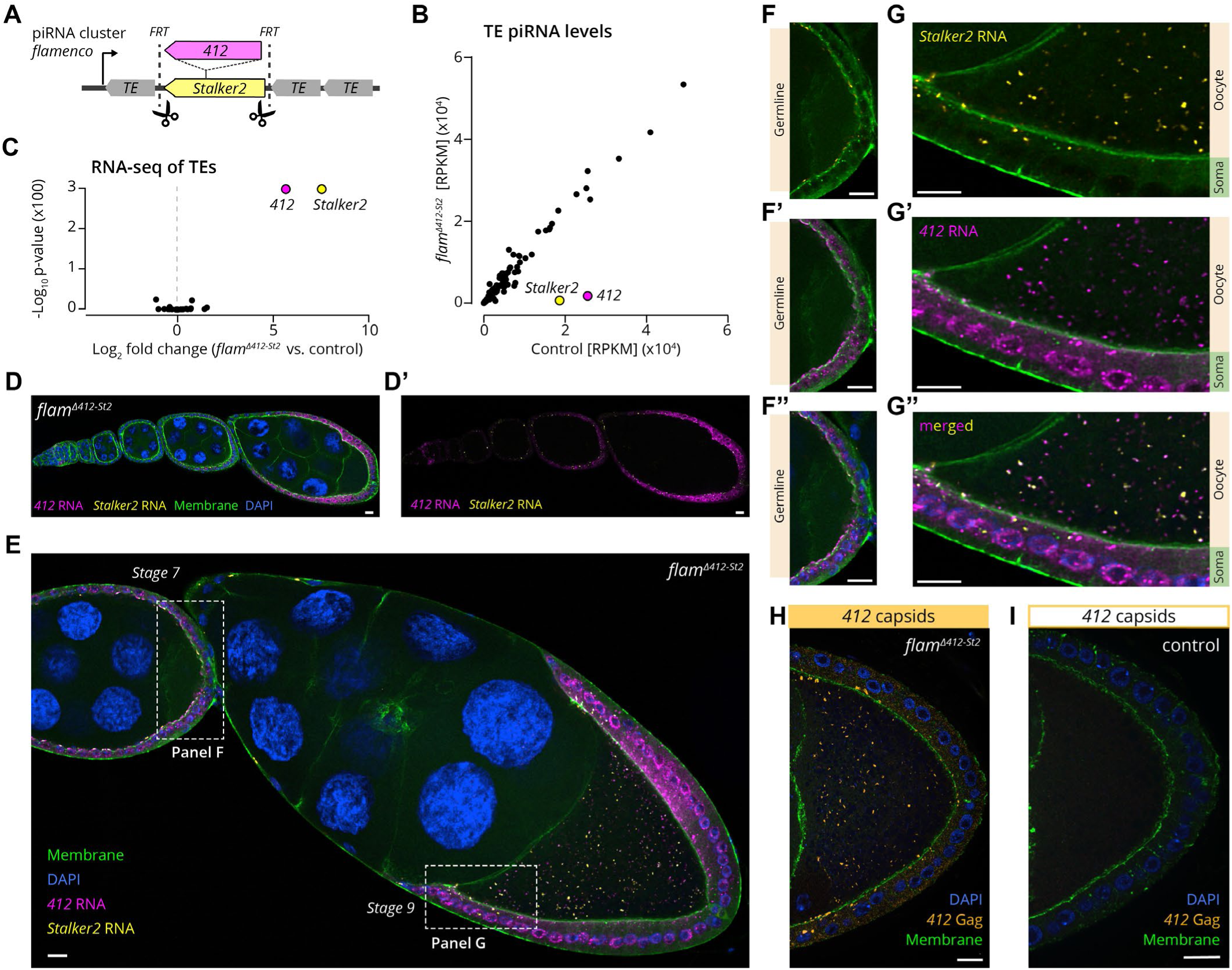
*412* and *Stalker2* retrotransposons autonomously infect the oocyte. **(A)** Schematic organization of the *flamenco* piRNA locus. The *flam^Δ412-St2^* allele was generated by removing a 15.5 kb region encompassing the *412-Stalker2* nested insertion. **(B)** Small RNA sequencing of piRNAs from *flam^Δ412-St^* ovaries, compared to control ovaries (identical genetic background). Each datapoint represents the number of antisense reads mapped to a TE consensus sequence, normalized as reads per kilobase of TE per million mapped microRNAs (RPKM). **(C)** Volcano plot of whole transcriptome poly(A)-enriched RNA-seq, from *flam^Δ412-St2^* ovaries, compared to control ovaries (n=3). Each datapoint represents the number of reads mapped to a TE consensus sequence. **(D-G)** Dual smFISH of *flam^Δ412-St2^* ovaries for *412* RNA (magenta) and *Stalker2* RNA (yellow) in **(D)** early developmental stages (1-7) **(E)** later developmental stages (7 and 9). **(F-G)** Zoomed-in regions from (E), showing absence of oocyte-localized signal for *412* and *Stalker2* in stage 7, whereas in (G) stage 9 there is smFISH signal for both *412* and *Stalker2* within the oocyte. **(H)** Whole-mount immunofluorescence of *flam^Δ412-St^* ovaries, showing somatic apical accumulation and oocyte localization of 412 Gag proteins. **(I)** Same as (H), but for control ovaries (identical genetic background). For (D-I), GFP-labelled myosin II (green) is used to demarcate both the oocyte and soma membranes. Scale bar: (D-I) 10 µm.

To investigate whether *412* and *Stalker2* are capable of autonomous cell-to-cell transmission, we tracked their RNA genome in *flam^Δ412-St2^*mutant ovaries using smFISH. As in the somatic piRNA pathway knockdown background, *412* and *Stalker2* transcripts were strongly expressed (Fig. 2D-E). During early oogenesis stages, transcripts of both elements showed distinct accumulations at the apical membrane of follicle cells (Fig. 2D-F). By stage 9, smFISH foci were also detected within the oocyte (Fig. 2E, G). We did not observe any smFISH signal in the cytoplasm of nurse cells, which are germline cells connected to the oocyte via ring canals (Fig. 2E). These findings suggest a specific cue or cellular organization that restricts *412* and *Stalker2* transfer exclusively to the oocyte, beginning mid-oogenesis.

To further test whether entire retroviral particles are transmitted, we analyzed the spatio-temporal distribution of potential 412 capsids using a specific polyclonal antibody targeting the CA (capsid) region of the Gag protein. Gag immunofluorescence signals were detected at the follicle cell membranes and within the oocyte, paralleling the RNA localization (Fig. 2H, I), which strongly suggests that entire *412* virus-like particles, encompassing the RNA genome inside a capsid, are capable of cell-to-cell transmission.

Altogether, the *412* and *Stalker2* retrotransposons are autonomous in their ability to transmit both their RNA genome as well as their capsid from somatic follicle cells into the oocyte and must employ a mechanism distinct from that of enveloped retroviruses.

### Two accessory small ORFs are expressed by MDG1 group retrotransposons

We hypothesized that members of the MDG1 group use an alternative infectivity factor to mediate cell-to-cell transmission. Two small open reading frames (ORFs) upstream of *gag*, denoted *sORF1* and *sORF2*, have been identified as putative accessory ORFs in the MDG1 group (Fig. 3A) (Avedisov et al., 1990; Costas et al., 2001; Makarova, 1997; Yuki et al., 1986). While their low non-synonymous/synonymous mutation rates suggest functionality (Costas et al., 2001; Mugnier et al., 2005), it remains unclear whether these short predicted ORFs (∼110 amino acids for sORF1 and ∼70 amino acids for sORF2) are indeed transcribed and translated into functional proteins and what their physiological roles might be.

**Figure 3:**
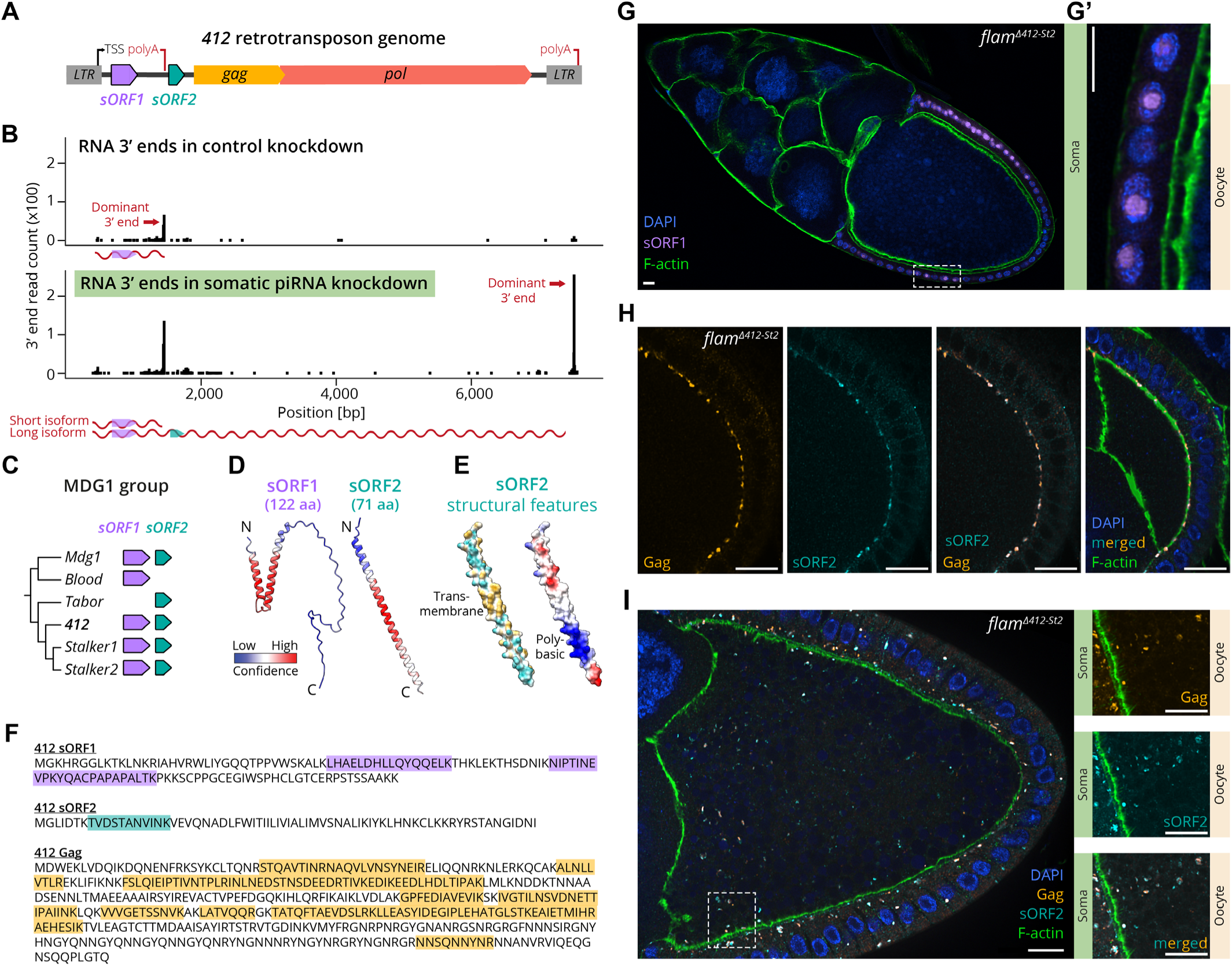
Characterization of the two unique sORFs, sORF1 and sORF2, encoded in *412* genome. **(A)** Schematic of the genomic features encoded by MDG1 retrotransposons, exemplified by the *412* consensus sequence. Features include flanking LTRs, capsid-encoding *gag* and enzymatic machinery-encoding *pol*. Two short ORFs (*sORF1* and *sORF2*), with a predicted premature transcription termination site (polyA) between them, are located upstream of *gag*. TSS, transcriptional start site. **(B)** Mapping of 3’ ends of long-read direct RNA reads corresponding to the *412* consensus sequence, in control ovaries (*Tj>Gal4, arr2-RNAi*, top) and somatic piRNA pathway knockdown ovaries (*Tj>Gal4, vret-RNAi*, bottom). The Y-axis indicates the number of reads, and the illustration represents the short and long isoforms produced based on the polyA site that is predominantly used. **(C)** Phylogenetic tree of the MDG1 clade based on *pol* alignment, with presence or absence of intact *sORF1* and *sORF2* sequences within the TE consensus sequences of *D. melanogaster* reference genome *iso-1*. Branch lengths are not to scale. **(D)** AlphaFold3 structural predictions for 412 sORF1 and sORF2, with cartoon representation color-coded by pLDDT confidence score (red: high, blue: low). **(E)** Notable features in sORF2 AlphaFold3 predicted structure: left, surface hydrophobicity (cyan: low, yellow: high); right, electrostatic surface potential (red: acidic, blue: basic). **(F)** Mass-spectrometry detection of peptides corresponding to sORF1 (purple), sORF2 (cyan) and Gag (orange) sequences in somatic piRNA pathway knockdown ovaries and OSCs. **(G)** Immunofluorescence staining of a stage 10 *flam^Δ412-St2^* follicle with α-sORF1 antibody (purple). (G’) Zoom-in of somatic nuclei in the boxed region in (G). **(H)** Co-localization of α-sORF2-N (cyan) and α-Gag (orange) on the apical somatic cell membranes in an early stage follicle. **(I)** Immunofluorescence signals of α-sORF2-C (cyan) and α-Gag (orange) often co-localize within the oocyte in later stage follicles. Right panels show zoomed-in region from I at a different Z-plane. For (G-I) F-actin is labelled with phalloidin (green) to demarcate cortical actin near the plasma membrane. Scale bars (G-I): 10 µm.

To this end, we performed long-read direct RNA sequencing on ovaries and a cultured cell line derived from ovarian somatic cells (OSCs), where in control conditions *Gypsy* retroviruses and MDG1 group retrotransposons are similarly repressed by the piRNA pathway. Upon somatic piRNA pathway knockdown, we detected continuous ∼7 kb transcripts of *412*, *Mdg1*, *Tabor*, *Stalker2* and *Blood* that span LTR to LTR, showing no evidence of splicing, and include the *sORF1* and *sORF2* region (Fig. 3B and fig. S3-4). Interestingly, an additional short ∼1.5 kb transcript terminating immediately after *sORF1* was also expressed, consistent with a previously predicted alternative polyadenylation site between *sORF1* and *sORF2* (Cherkassova et al., 1991). This short transcript could be detected at low levels for most MDG1 elements also in the control ovaries and OSCs. Our data therefore suggest that in wildtype conditions, a short *sORF1* transcript is expressed, while full-length transcripts spanning both *sORFs* together with *gag* and *pol* are repressed by the piRNA pathway.

Based on Berkeley *Drosophila* Genome Project (BDGP) and Repbase TE consensus sequences, *sORF1* and *sORF2* genes are encoded in all members of the *D. melanogaster* MDG1 group, except for *Tabor* and *Blood*, which harbor deletions disrupting *sORF1* and *sORF2*, respectively (Fig. 3C) (Bao et al., 2015; Costas et al., 2001; Kaminker et al., 2002). Protein structure predictions using AlphaFold3 (Abramson et al., 2024) revealed a potential helix-turn-helix motif for sORF1, while sORF2 likely forms an alpha-helix with a central hydrophobic domain followed by an adjacent polybasic region (Fig. 3D-E). To determine whether the two sORFs are translated, we conducted untargeted proteomics on somatic piRNA-deficient ovaries and OSCs. We identified high confidence peptides corresponding to sORF1 for *412*, *Mdg1*, *Blood* and *Stalker2* and to sORF2 for *412*, *Mdg1* and *Stalker1* (Fig. 3F, fig. S5. We note that the detection of sORF2 is challenging due to the limited number of suitable tryptic peptides).

To gain insight into their potential functions, we determined the cellular localization of the *412* sORF1 and sORF2 proteins in ovaries using polyclonal antibodies. Antibody specificity was validated by the strongly elevated signal in *flam^Δ412-St2^* ovaries versus control ovaries from flies that were genetically identical except for the targeted *flamenco* deletion (fig. S6A-D). Whole mount immunofluorescence experiments revealed that sORF1 is exclusively expressed in somatic follicle cells, where it localizes to a subnuclear compartment (Fig. 3G). No signal was detected in the oocyte, even at later developmental stages when *412* transmission occurs. We conclude that sORF1 likely functions early in the transposon life cycle and is not directly involved in the transmission process.

In contrast, two independent antibodies targeting the N or C-terminus of *412* sORF2 revealed a plasma membrane-corresponding staining in follicle cells (Fig. 3H and fig. S6B, E). We obtained similar results in flies expressing a C-terminally HA-tagged sORF2 transgene in the soma, where the HA antibody and the sORF2 N-terminal antibody yielded overlapping signals (fig. S6C). During previtellogenic stages (up to stage 6), sORF2 localized to the apical side of follicle cells, which faces the germline (fig. S6E). Notably, these membranal sORF2 accumulations were also enriched with viral capsids, as indicated by co-localization with the Gag-targeting antibody and with cortical actin marker phalloidin (Fig. 3H). At later developmental stages (from stage 7 onwards), we detected abundant sORF2 signal also within the oocyte, often overlapping with Gag, in accordance with the timing of *412* transmission into the oocyte observed by smFISH (Fig. 3I).

In sum, the plasma membrane localization of sORF2 and its close association with viral capsids suggest that sORF2 may play a role in facilitating the cell-to-cell transmission of MDG1 group retrotransposons.

### sORF2 is a transmembrane protein with hallmarks of cell-cell fusogens

To gain insights into sORF2 function, we examined conserved features in MDG1 group sORF2 proteins using a multiple sequence alignment (Fig. 4A). This analysis revealed a highly hydrophobic central domain predicted to function as transmembrane domain with high-confidence, flanked by an N-terminal ectodomain and a C-terminal cytoplasmic tail (see Materials and Methods). Furthermore, all sORF2s harbor an amphipathic motif in their ectodomain and a positively charged basic patch followed by an amphipathic motif in their cytoplasmic tail (fig. S7). Finally, four out of six sORF2 proteins harbor a predicted N-terminal glycine myristoylation motif (MGxxxS/T), a fatty acid modification known to promote protein-membrane association and localization to specific membrane microdomains, especially in the context of viral proteins (Wang et al., 2021).

**Figure 4:**
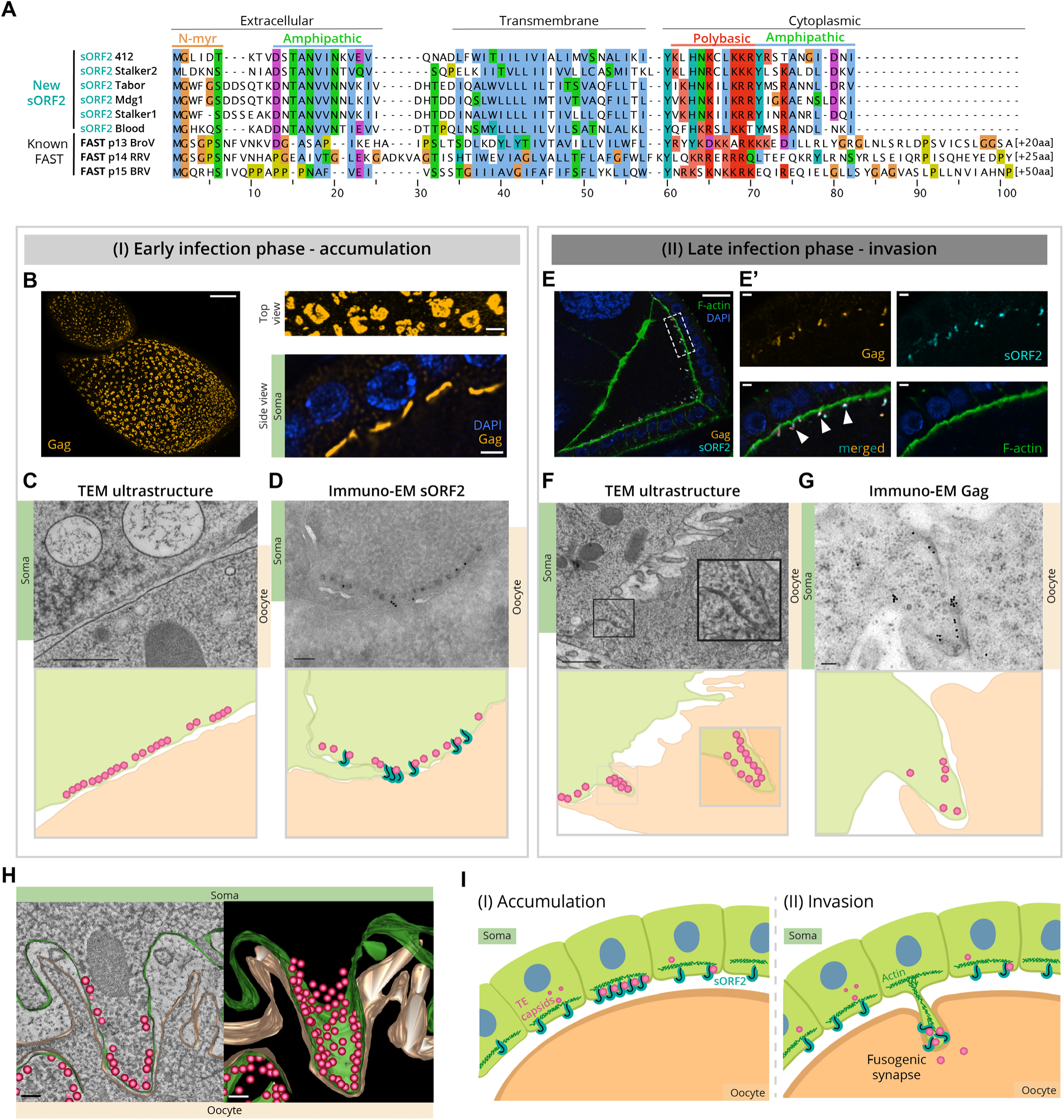
sORF2 proteins resemble FAST proteins, and promote cell fusion by facilitating proximity of membranes. **(A)** Multiple sequence alignment of sORF2 proteins from *D. melanogaster* and three previously described FAST proteins. p13 from bat Broome reovirus (BroV, ACU68609.1); p14 from reptilian orthoreovirus (RRV, AAP03134.1); p15 from baboon orthoreovirus (BRV, AAL01373.1). sORF2 proteins of *Stalker1* and *Stalker2* lack the predicted canonical N-myristoylation motif (N-myr, MGxxxS/T) but share all other features. *Blood* consensus sequence lacks an intact sORF2, but several heterochromatic insertions supported a *Blood* variant with the presented sORF2 sequence. Clustal X coloring scheme labels conserved residues according to amino acid profile, additional Arg and Lys residues in the polybasic region are shown in light red. **(B)** Left, Immunofluorescence Z-stack projection of stage 7 *flam^Δ412-St2^* follicles expressing GFP-labelled myosin II (green)stained with α-Gag (orange) showing accumulations of capsids within the somatic follicle cell layer covering the oocyte. Scale bar: 10 µm. Top right, zoom-in of the capsid accumulation pattern. Scale bar: 2 µm. Bottom right, mid-section of the follicle showing 412 capsid accumulations along the somatic apical membranes. Scale bar: 2 µm. **(For C-D, F-G)** Upper panels, Transmission (TEM) and immuno-EM images. Lower panels, schematic representation of EM images, somatic cells (green) with observed capsids (pink) are in close proximity to the oocyte membrane (beige) and sORF2 proteins (cyan). **(C)** TEM of capsid accumulations along somatic apical membranes in stage 7 *flam^Δ412-St2^* follicle. Scale bar: 500 nm. **(D)** Immuno-EM of *flam^Δ412-St2^* stage 7 follicle using α-sORF2-C antibodies, labelled with secondary antibody-conjugated gold particles (black dots in upper panel, cyan in lower schematic), revealing sORF2 localization near accumulated capsids. Scale bar: 100 nm. **(E)** Immunofluorescence of *flam^Δ412-St2^* ovaries co-stained with α-Gag (orange), α-sORF2-C (cyan) and phalloidin (green), showing co-localization in protrusions extending into the oocyte (white arrowheads). Scale bars: 10 µm, zoom-in panels on right: 2 µm. **(F)** TEM of an invasive protrusion filled with capsids in *flam^Δ412-St2^* follicle, extending into the oocyte. Adjacent physiologically normal spacing between membranes and typical microvilli are shown for comparison. Scale bar: 500 nm. Inset, zoom-in of protrusion with indistinguishable soma and oocyte membrane. **(G)** Immuno-EM of *flam^Δ412-St2^* ovaries using α-Gag antibody, with gold particles (black dots) corresponding to 412 capsids within an invading protrusion. Scale bar: 100 nm. **(H)** Snapshot of electron tomography segmentation and 3D rendering of an invasive protrusion in *flam^Δ412-St2^* ovaries, filled with 412 and Stalker2 viral-like particles (pink). The somatic membrane (green) is closely associated with the oocyte membrane (beige), with minimal separation between them. Scale bar: 100 nm. **(I)** Proposed model for soma-to-oocyte infectivity of MDG1 LTR retrotransposons. (Left) somatically expressed transmembrane sORF2 proteins (cyan) are closely associated with LTR retrotransposon capsids (pink). Upon local clustering and additional unknown cues, cortical actin-remodeling (green) is initiated (right), promoting the formation of invasive protrusions toward the oocyte membrane. Once the membranes are close enough in a fusogenic synapse structure, transient fusion between somatic and oocyte membranes enables LTR retrotransposons to enter the oocyte.

Collectively, these structural features are highly reminiscent of fusion-associated small transmembrane (FAST) proteins, which mediate cell-cell fusion rather than classical viral-cell fusion (Duncan, 2019). FAST proteins have been mainly identified in several fusogenic reoviruses and rotaviruses (order *Reovirales*), which are non-enveloped, double-stranded RNA viruses that infect a wide range of hosts (Ciechonska & Duncan, 2014; Diller et al., 2019). Recent studies have shown that FAST proteins function as cell-cell fusogens by multimerizing in specific membrane domains and recruiting the actin cytoskeleton to generate localized mechanical force, pushing two cellular membranes together and disrupting them to drive membrane fusion (Chan et al., 2020, 2021). Since FAST proteins are short (∼100-200 residues) and exhibit highly divergent sequences, primary sequence-based homology search cannot be used to establish a link between sORF2 and FAST proteins. Nevertheless, the MDG1 group sORF2 and reoviral FAST proteins share a striking resemblance of domain organization and multiple predicted motifs (Fig. 4A).

### 412 sORF2 and capsids localize to fusogenic synapses between follicle cell protrusions and the oocyte membrane

In the *Drosophila* egg chamber, somatic follicle cells form apical microvilli - actin-filled membranous extensions characteristic of epithelial cells - that project toward the oocyte (Schlichting et al., 2006). We wondered whether MDG1 group sORF2 proteins function in an analogous manner to reoviral FAST proteins, by forcing somatic microvilli membranes into close contact with the oocyte membrane to enable local cell-cell fusion and capsid transmission. To test this hypothesis, we performed extensive light and electron microscopy (EM) studies, leveraging the natural contrast provided by viral capsids, to examine the follicle cell-oocyte interface in *flam^Δ412-St2^*and control ovaries.

Using EM, we observed that the soma-to-oocyte infection process occurs in two distinct phases: In the first phase, 412 virus-like particles and sORF2 proteins accumulate in discrete clusters along the apical membranes of somatic follicle cells, in agreement with our immunofluorescence findings (Fig. 3H, 4B-C). Transmission EM ultrastructure analysis revealed that the abundant capsid structures measured 42+/-5 nm in diameter and exhibited a characteristic dark outer rim and a lighter inner region, consistent with the size and pattern reported previously for *Gypsy* retroviruses (Lécher et al., 1997). Cryo-immunoEM experiments with antibodies targeting 412 Gag and sORF2 confirmed that many of the observed virus-like particles correspond to 412 capsids and closely associated sORF2 proteins. However, given the co-expression of *Stalker2*, we anticipate the virus-like particles to represent a mix of 412 and Stalker2 capsids (Fig. 4D, fig. S8).

The second phase is characterized by the formation of invasive protrusions at the apical membrane of follicle cells which extend into the oocyte (Fig. 4E-H, fig. S8C). These cellular protrusions were exclusively observed within the ultrastructure of *flam^Δ412-St2^* ovaries and typically had enlarged bulges at their tips, each filled with dozens of capsids (Fig. 4F-G). Unlike regular microvilli, which are sleek, slender and devoid of capsids, these specialized structures were distinguishable by their bulging morphology and tight encapsulation by the oocyte membrane (fig. S8F-G). EM tomography experiments further revealed that capsids are dispersed along the periphery of the invasive protrusion bulges, reinforcing their role as functional structures for viral transmission (Fig. 4H and Supplementary Movie S1).

In stage 6-7 *flam^Δ412-St2^* follicles, the invasive protrusions displayed considerable variation in size, shape, position, and number (fig. S8F-G, S9A-B), likely reflecting different maturation stages in the soma-to-oocyte infection process. Interestingly, these protrusions were rarely observed at earlier (stage 4–5) or later (stage 9–10) developmental stages, suggesting a narrow window of opportunity for soma-to-oocyte transfer. This window corresponds to the time from the onset of microvilli formation to the stage when the secreted vitelline bodies coalesce to form the impenetrable vitelline shell around the oocyte (fig. S9C-D) (King & Koch, 1963; Mallart et al., 2024). We never observed the characteristic protrusions in somatic cells neighboring the nurse cells, although capsids readily accumulated along the apical membrane, corresponding to the first phase only (Fig. 4B, fig. S9E-F). These findings imply the critical involvement of unidentified tissue-specific host factors in the formation of the invasive protrusions.

In order to retain the native ultrastructure of membranes, we used high pressure freezing for the EM ultrastructure experiments, allowing the clear visualization of the lipid bilayers of somatic follicle cell and oocyte membranes. Intriguingly, in several capsid-filled protrusions, we observed that the somatic and the oocyte membranes were indistinguishable, suggesting sites of confined, intimate membrane-membrane interaction that overcame the repulsive forces acting between membranes (Chernomordik & Kozlov, 2003) (notice spacing between membranes in Fig. 4F,H and in fig. S8,9). We propose that these sites represent restricted zones of transient cell-cell fusion, facilitating the transmission of viral-like particles into the large oocyte volume (Fig. 4I).

Based on our detailed cell-biological studies, we suggest that the invasive protrusions function as fusogenic synapses, specialized structures that mediate localized membrane proximity and fusion between adjacent cells (J. H. Kim & Chen, 2019; Shilagardi et al., 2013). These findings strongly support a model in which sORF2 drives the formation of these protrusions, similar to FAST proteins, enabling MDG1 group retrotransposons to transmit directly from somatic follicle cells into the oocyte.

### sORF2-encoding LTR retrotransposons are widespread in insect genomes

Many organisms share the general architecture of ovarian follicles, where supportive somatic cells and their membrane extensions closely interact with the oocyte (Clarke, 2022; Jaglarz et al., 2008). To explore whether the sORF2-mediated transfer demonstrated in *Drosophila* represents a broader phenomenon, we examined the prevalence of sORF2/FAST-like proteins in LTR retrotransposons (Fig. 5A). We first performed a non-exhaustive search on ∼8,500 representative *Metaviridae* consensus sequences in arthropods taken from the Repbase database (Bao et al., 2015). Due to the short length of sORF2 proteins, direct sequence-based homology searches were unproductive. Instead, we translated all possible ORFs from the consensus sequences and filtered for short ORFs harboring a predicted transmembrane domain and N-myristoylation motif (Fig. 5B-C, see Materials and Methods). This approach identified 105 high-confidence hits across 29 organisms, ranging from flies and mosquitoes (*Diptera*), through butterflies and moths (*Lepidoptera*) to beetles (*Coleoptera*), (Fig. 5B). The highest diversity of sORF2/FAST-like sequences within a single organism, 33, was found in the mosquito *Aedes aegypti*, the primary vector of human-pathogenic viruses including dengue, Zika, and yellow fever viruses, which has an exceptionally large, repeat-rich genome (Daron et al., 2024). All *sORF2/FAST*-like candidates were found in the forward orientation relative to the *gag* and *pol* ORFs, suggesting they depend on the primary transcriptional start site and do not utilize an alternative promoter (fig. S10A). In some LTR retrotransposons of *Lepidoptera*, we have identified cases where *sORF2* genes are encoded downstream of *pol* instead of upstream of *gag* (7/22 consensus sequences in *Lepidoptera*).

**Figure 5:**
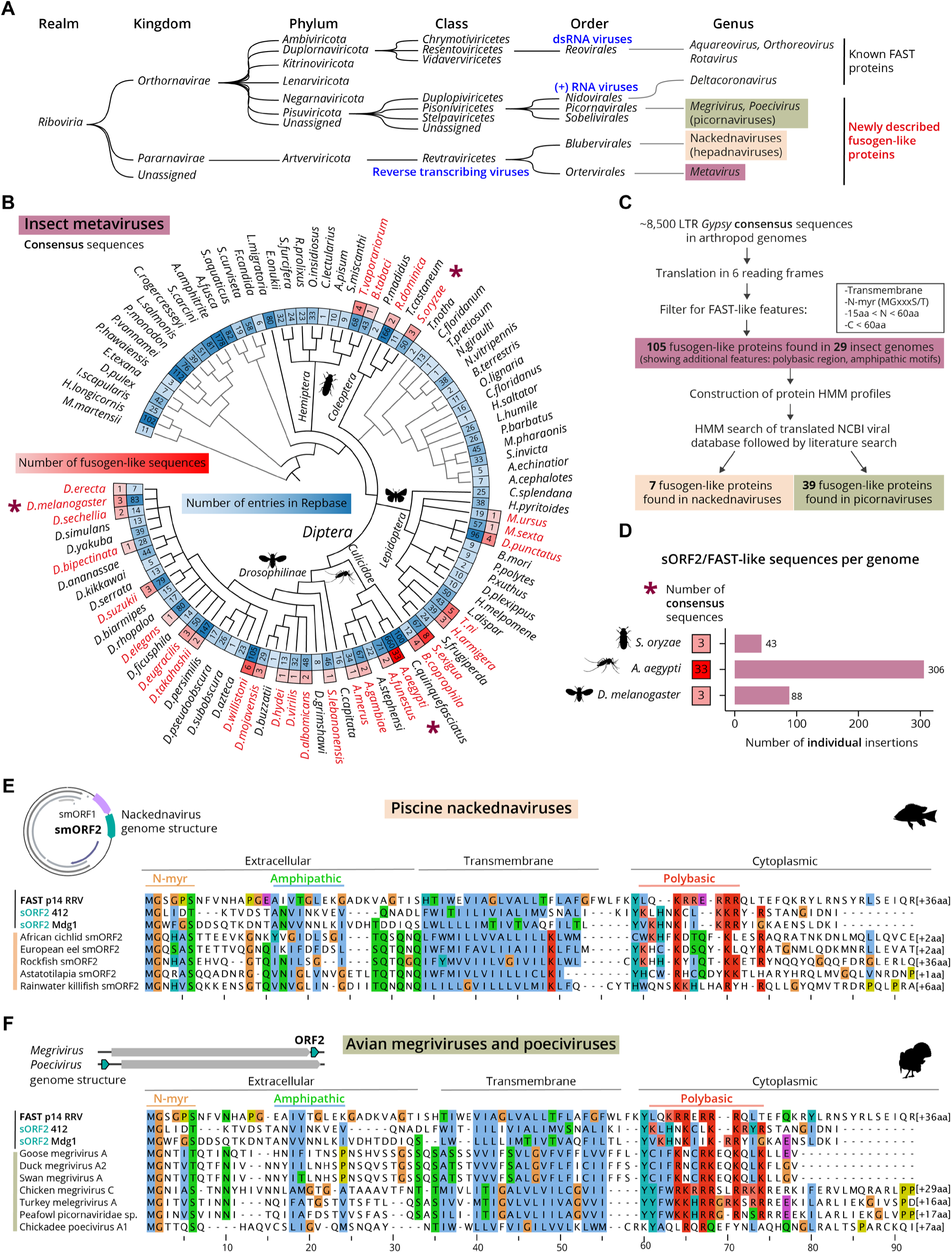
sORF2/FAST-like sequences are abundant and widespread among non-enveloped viruses. **(A)** Taxonomic tree of all viruses and TEs described in this study. **(B)** Taxonomic tree of arthropod genomes annotated in Repbase for *Metaviridae* consensus sequences. Inner circle (blue shades) – number of TE entries in the database per species. Outer circle (red shades) – number of *sORF2/FAST*-like consensus sequences found per species. Insect genomes analyzed in (D) are labelled with asterisk. **(C)** Schematic representation of computational pipeline to identify sORF2/FAST-like proteins in additional genomes. **(D)** Number of individual sORF2/FAST-like sequences in LTR retrotransposon insertions of three insect genomes. **(E-F)** Multiple sequence alignment of a subset of sORF2/FAST-like (E) smORF2 proteins found in fish-infecting nackednaviruses (F) ORF2 proteins from bird-infecting picornaviruses, and their typical genome organization. Clustal X coloring scheme labels conserved residues according to amino acid profile, additional Arg and Lys in the polybasic region are shown in light red.

A closer examination of the multiple sequence alignment of the 105 high-confidence sORF2/FAST-like sequences revealed additional conserved features previously described for FAST proteins, including a juxtamembrane polybasic region and amphipathic motifs (fig. S7C), despite these not being part of the initial search criteria. sORF2/FAST-like sequences ranged in length from 63 to 107 amino acids, with shorter variants usually found in flies and mosquitoes, and longer ones in butterflies, moths, and beetles. By comparison, known FAST proteins in *Reoviridae* range from 80 to 200 residues (Duncan, 2019; Veletanlic et al., 2023). Interestingly, in both known FAST proteins and newly found sORF2-like proteins, the primary variation in overall length arises from differences in the endodomain, while the ectodomain is rather constant at around 30 residues (fig. S10B). This conserved ectodomain length may reflect a functional role in membrane biology and could serve as a useful narrowing filter for identifying additional *bona fide* sORF2/FAST-like proteins. Finally, a previously undescribed feature that has emerged with the increased number of available sequences is a conserved Tyr residue immediately downstream of the transmembrane domain. This residue is often found at the interfacial boundary of transmembrane proteins, forming part of the “aromatic belt” (Baker et al., 2017) (fig. S10B).

Estimating the prevalence of *sORF2/FAST*-like genes within a genome is challenging because consensus sequences fail to capture the full extent of heterogeneous multi-copy TE insertions. To overcome this limitation, we analyzed three high-quality TE-curated insect genomes, systematically scanning them for sORF2/FAST-like sequences within LTR retrotransposon insertions. Using this approach, we identified 88 insertions in the *Drosophila melanogaster iso-*1 reference genome, mostly belonging to *412*, *Mdg1*, *Tabor* and *Blood* retrotransposons (Fig. 5D). In the mosquito *A. aegypti*, 306 LTR retrotransposon insertions with *sORF2/FAST*-like sequences, of which 296 corresponded to the 33 representative consensus sequences, were found. Lastly, in the major agricultural pest *Sitophilus oryzae* (rice weevil), 43 *sORF2/FAST*-like sequences were found in the context of LTR retrotransposon insertions. Altogether, considering the enormous diversity of insect species and the abundance of sORF2/FAST-encoding LTR retrotransposons within them, our findings highlight a highly successful viral-like evolutionary strategy of LTR retrotransposons to prevail within their hosts. Further supporting this notion is the evidence that the *412* retrotransposon invaded *D. melanogaster* genome less than 200 years ago, and has since successfully spread to every sampled natural population of *D. melanogaster* (Scarpa et al., 2024).

### sORF2/FAST-like proteins suggest an Envelope-independent transmission strategy also in hepadnaviruses and picornaviruses

If FAST and sORF2-like proteins enable cell-to-cell transmission in non-enveloped fusogenic reoviruses and LTR retrotransposons, respectively, we hypothesized that other non-enveloped viruses might employ similar strategies. Building on the large repertoire of sORF2/FAST-like proteins identified in insect genomes, we constructed hidden Markov model (HMM) profiles of sORF2s and used them to scan the NCBI viral database. This approach revealed putative sORF2/FAST-like proteins in two additional, entirely unrelated viral families: *Hepadnaviridae* and *Picornaviridae*.

Hepadnaviruses are considered distant relatives of retroviruses (Fig. 5A) (Magnius et al., 2020). They also undergo reverse transcription as part of their life cycle and encode for an Envelope protein, although distinct from that of retroviruses. The most extensively studied member of this group is human hepatitis B virus, whose phylogenetic origin remains unresolved. In an effort to elucidate the time and source of *envelope* acquisition, Lauber *et al*. described a family of non-enveloped hepadnaviruses in teleost fish, which they named “nackednaviruses” (Lauber et al., 2017). These fish viruses exhibit many canonical structural and functional features of hepatitis B virus, except for the *envelope* gene, leaving the mechanism of how these viruses enter cells unclear.

Interestingly, the authors note that all thirteen described nackednaviruses encode two small ORFs, designated “*smORF1*” and “*smORF2*”, positioned in the viral genome in the same orientation and upstream of the capsid-encoding core gene (*orf C*). We propose *smORF2* encodes for a sORF2/FAST-like protein, as all eleven complete smORF2 proteins are 75-114 amino acids long, contain a predicted transmembrane domain with N-outside/C-inside topology, and exhibit amphipathic motifs in both the extracellular and cytoplasmic domains. Additionally, they feature a polybasic region just C-terminal of the membrane domain, and seven encode a canonical N-myristoylation motif (Fig. 5E, fig. S11A). Available nackednaviral RNA-derived sequences indicate that both *smORFs* are transcribed and not spliced out. Based on a later study of the nackednavirus capsid, no structural change of the capsid occurs upon pH change, in contrast to capsids of known non-enveloped viruses, which would argue against a typical endocytic-entry path (Pfister et al., 2023). We thus speculate that nackednaviruses employ a cell-to-cell transmission mode involving their sORF2/FAST-like smORF2 proteins. Direct experimental studies are required to determine the viral and cellular determinants of smORF2 activity.

The second group that was found using the sensitive HMM search was avian picornaviruses, specifically megriviruses and poeciviruses. Unlike most picornaviruses, which encode a single polyprotein, some megriviruses encode an accessory protein, ORF2, of unknown function (Boros et al., 2014). We found many of the characteristic features of FAST proteins as described above in 34 megriviruses sequences, at a conserved position downstream to the main ORF, spanning 67-115 amino acids in length. (Fig. 5F, fig. S11B). Interestingly, pathological reports of turkey hepatitis cases associated with megrivirus infections describe syncytia-like cells in the liver, which the authors note is peculiar since there is no established relationship between syncytia and a picornavirus (Hauck et al., 2014). We suggest the syncytia were formed by the activity of sORF2/FAST-like ORF2 proteins encoded in the megrivirus genome, which is circumstantially lacking from other non-syncytia forming picornaviruses. Similarly, the related poecivirus, the suspected causative agent of the devastating avian keratin disorder in birds, also encodes an unannotated gene for a sORF2/FAST-like protein, however it is positioned upstream of the main ORF (Fig. 5F, fig. S11B) (Zylberberg et al., 2018). Future work is needed to determine what is the evolutionary trajectory and possible shared origin of sORF2/FAST-like proteins found in reoviruses, rotaviruses, LTR retrotransposons, hepadnaviruses, and picornaviruses.

## DISCUSSION

This study provides direct evidence for viral-like infectivity of MDG1 group LTR retrotransposons in *Drosophila*, demonstrated at both the RNA and protein levels. We show that the soma-to-germline transmission occurs independently of Envelope and relies instead on the formation of somatic protrusions that invade the oocyte. In the context of the host germline, MDG1 group retrotransposons are yet another example of breaking the Weismann’s barrier in metazoans, which postulates genetic material is only unidirectionally transferred from germline to soma (Bline et al., 2020; Yoth et al., 2023).

Direct cell-to-cell transmission offers several advantages over Envelope-mediated cell exit and re-entry, such as avoiding extracellular restriction factors and bypassing the need for receptor-mediated internalization. While cell-cell fusion has been described for some retroviruses, such as HIV and the case of conduits formed by HTLV (Bracq et al., 2018; Van Prooyen et al., 2010), these processes are thought to depend on Envelope proteins. Our work demonstrates, for the first time, a functional *env*-independent retrovirus capable of direct transmission.

The sORF2 protein, bearing structural similarity to reoviral FAST proteins—the only known fusogenic proteins encoded by non-enveloped viruses—appears to enable limited and specific cell-cell fusion. Unlike the aggressive fusion seen in fusogenic reoviruses, which leads to extensive syncytium formation, sORF2-mediated fusion appears to be tightly controlled. This regulation likely ensures host fertility while enabling retrotransposon persistence. The mechanism of sORF2-mediated fusion resembles more developmental cell fusogens, such as Myomerger and EFF-1, rather than potent viral fusogens (Sapir et al., 2008). We propose that sORF2 evolved to be temporally and spatially restricted, allowing viral genome transfer only in a subset of cells in a well-defined developmental window. Invasive protruding membrane structures, as seen in fertilization (Satouh & Inoue, 2022), osteoclast fusion (Oikawa et al., 2012), macrophage fusion (Faust et al., 2019), and myoblast fusion (Luo et al., 2022; Shilagardi et al., 2013) are distinctive features of fusion-related processes *in vivo*. However, such structures associated with FAST proteins have not yet been observed in their native tissue context. Our study synthesizes these findings together, suggesting that in the ovary tissue, FAST-like sORF2 proteins either exploit existing actin-rich microvilli or form them *de novo* to bring the somatic and oocyte membranes closely together in a fusogenic synapse structure.

To the best of our knowledge, less than 20 FAST proteins have been reported, mostly within the *Reoviridae* family (Duncan, 2019; Veletanlic et al., 2023) and a few others in coronaviruses (Huang et al., 2016; Sartalamacchia et al., 2024). Our findings reveal that sORF2/FAST-like proteins are encoded also in *Metaviridae*, *Hepadnaviridae* and *Picornaviridae*, substantially expanding the repertoire of known FAST-like proteins across the viral kingdom. Given the multi-copy nature of TEs and the extraordinary diversity of insects, birds, and fish, as well as their short and non-conserved primary amino acids sequence, these proteins are likely far more abundant than previously appreciated. Among these different virus-host interactions, perhaps the most surprising is that of insect LTR retrotransposons, which were thought to be confined to the cells in which they are expressed.

Viruses are known for their ingenious strategies to manipulate host cell biology, often inspiring biotechnological advances. Similar to FAST proteins, which have recently shown promise in gene delivery systems (Brown et al., 2024), sORF2-based mechanisms could inform the development of targeted biotechnological tools or therapeutic strategies. Furthermore, studying host-pathogen interactions in insects may yield valuable insights for controlling mosquito-borne diseases or developing biocontrol agents against agricultural pests. Our work provides fundamental new insights on retrotransposon biology, fusogen evolution, and host-pathogen interactions, with implications for both fundamental and applied sciences.

### Limitations of the study

Despite these advances, our study has some limitations. First, our efforts to recapitulate sORF2-mediated fusion in minimal heterologous experimental systems using standard cell-cell fusion reporter assays have not proven successful so far, likely due to specific requirements for cellular factors and tissue-level architecture that is challenging to replicate in cell culture. Another confounding factor in our study is the encoding of sORF2 as part of the multi-copy transcript of MDG1 group elements, which hinders genetic approaches to determine if sORF2 is indeed required for cell-cell fusion. Lastly, the identification of sORF2/FAST-like proteins remains constrained by our search criteria, which focus on canonical start codons and N-terminal myristoylation motifs, excluding non-canonical or alternative variants, such as palmitoylated proteins seen in some FAST proteins (Duncan, 2019; Shmulevitz et al., 2003). Future studies need to address these limitations to deepen our understanding of sORF2 function and its evolutionary origin.

## Supporting information

Supplemental Table S1: sORF2/FAST-like sequences identified in this study

Supplementary Table S2: Fly genotypes, smFISH probes and siRNA sequences used in this study

Data S2: High resolution oocyte-soma interface in control follicle

## ACKNOWLEDGEMENTS

We thank the Vienna BioCenter Core Facilities (VBCF) for extensive technical support, knowledge and expertise: the BioOptics facility for imaging experiments, the Fly&Worm facility for transgenesis and CRISPR-mediated genome engineering, the NGS facility for RNA sequencing experiments, the Proteomics facility, particularly Karl Mechtler and Elisabeth Roitinger, for mass-spec experiments, the Protein Chemistry facility for assisting in raising the polyclonal antibodies, as well as the Electron Microscopy facility in which EM and ET experiments were performed. Proteomics analyses were performed by the Proteomics Facility at IMP/IMBA/GMI using the VBCF instrument pool. We thank Laszlo Tirian and Jakob Schnabl for technical support, Emilie Brasset for sharing the *412* anti-Gag antibody, as well as Josquin Daron and Louis Lambrechts for the *A. aegypti* TE library. We thank the Bloomington and VDRC stock centers for flies. We thank Jackson Ridges, Victoria Deneke, Andrea Pauli, Alejandro Burga, Aude Bernheim, Charlotte Mallart and past and current members of the Brennecke lab for fruitful discussions and for comments on the manuscript.

## AUTHOR CONTRIBUTIONS

**MV**: Conceptualization, methodology and analysis of all experiments unless stated otherwise, writing. **AB**: EM experiments methodology and analysis. **MN**: Computational analysis, software and data curation. **DH:** Methodology and computational analysis of ONT and Illumina RNA-seq, small RNA-seq, and investigation of TE distribution. **PM**: Technical support in methodology and validation. **BR**: Methodology and analysis of LTR reporters. **PD**: Methodology of *flam^Δ412-St2^* and support of fly genetics. **KAS**: Conceptualization, phylogenetic investigation, methodology and analysis of RNA-seq and smFISH in somatic piRNA pathway knockdown experiments, resources and extensive support in fly genetics, funding. **JB**: Conceptualization, analysis, writing, supervision, funding and infrastructure.

## DATA AND CODE AVAILABILITY

All fly stocks and antibodies generated for this study are available upon request. The sequencing data have been deposited in the NCBI Gene Expression Omnibus (GEO) under the accession number XXX. The mass spectrometry proteomics data have been deposited to the ProteomeXchange Consortium via the PRIDE (Perez-Riverol et al., 2025) partner repository with the dataset identifier PXD061763.

## Materials and Methods

### Flies

#### Drosophila melanogaster husbandry and strains

Flies were grown at 25 °C with 12h dark/light cycles under standard laboratory conditions. For dissections of ovaries, 1-3 days old flies post-eclosion were transferred to cages with daily changed apple juice plates and yeast paste for 2 days. Ovaries were collected and stored in ice-cold PBS until fixation for up to 30 minutes after dissection. All fly genotypes are described in Supplementary table 2.

#### Somatic piRNA pathway knockdown flies

Tissue-specific knockdown of the piRNA/PIWI pathway was obtained by crossing *traffic jam*-*GAL4*, to UAS-*vreteno*^GD^ flies (somatic piRNA pathway knockdown) or UAS-*arrestin2^GD^* (control knockdown) (Dietzl et al., 2007; Olivieri et al., 2010). *Tj-GAL4* was recombined with *UAS-myrGFP* (myristoylated GFP) to label somatic membranes in smFISH and in immunofluorescence experiments.

#### Generation of *flam^Δ412-St2^* flies

To delete the single full-length insertion of *412* from the *flamenco* locus, gRNAs targeting unique regions upstream (ggatctatttcctggacac) and downstream (ggtggcttcacaaaacacga) of *412-Stalker2* were cloned into a pDCC6b plasmid (Gokcezade et al., 2014) and co-injected with an HDR donor oligo (IDT) to introduce FRT sites into embryos containing the *iso-1* X-chromosome. Successful targeting events were identified by PCR and confirmed by Sanger sequencing.

<u>FRT donor upstream sequence:</u> ggtccaaaaaccttctagcttgccctctggaccaaactggatctatttcctgGAAGTTCCTATaCtttctagaGAATAGGAACTTCgGAATAGGAACTTCacaggaccaaagtcgcgcgcgttctcacaactcgtatttagttttcgcaatctacc

<u>FRT donor downstream sequence:</u> ggtatgttaagtttataatattttacgccaatttccgcaagccggtggcttcacaaaacaGAAGTTCCTATTCcGAAGTTCCTATTCtctagaaaGtATAGGAACTTCgacggagtaacttttaagaactctttattgagtagagcaagtgctgttgcttatgagg

We generated female flies heterozygous for both FRT insertions in *trans* and carrying a *nanos-flp* construct on the 2^nd^ chromosome (Kaushal et al., 2021). This resulted in the FRT-Flp mediated deletion of a 15.5kb region between the two FRT sites in the progeny of those females. Individual offspring were screened by PCR for this deletion and verified by Sanger sequencing. Two independent lines (#1 and #2) were retained for further experiments. Control flies (*FRT*) were generated by retaining the integrated FRT site without the Flp recombination, representing genetically identical flies except the *flamenco* deletion. Both *flam^Δ412-St2^* and the control flies were crossed to *iso-1/CyO* flies to isogenize the 2^nd^ chromosome (*flam^Δ412-St2^*;*iso-1/iso-1*;;) and to GFP-labelled Myosin II (*zip*) to label membranes.

#### Generation of *flam^Δ412-St2^*; p(*flam*)-PR412sORF2-HA flies

A piRNA-resistant *412 sORF2* (PR412sORF2) sequence was designed by altering wobble positions of the original *412 sORF2* consensus sequence, and synthesized as a G-block (IDT). The PR412sORF2 was tagged with a short linker and 3xHA at the C-terminal and cloned into the *flamenco* reporter vector (Senti et al., 2023) by replacing the *lacZ* ORF, downstream of a putative ∼4 kb *flamenco* promoter to drive constitutive expression in somatic follicle cells. The construct was integrated into *attp40* and balanced flies were subsequently crossed to *flam^Δ412-St2^* and *FRT* control flies.

#### LacZ LTR reporter transgene and functional analysis

The 514 bp LTR of *412* was cloned upstream of a *lacZ* gene and integrated into *attP40* as in (Senti et al., 2023). The constructs were combined with piRNA pathway knockdown RNAi lines *UAS-vreteno^GD^*and *UAS-arrestin2^GD^* for somatic piRNA pathway knockdown, or with shRNA-lines targeting *aub+ago3* or *white* (control) for germline piRNA pathway knockdown. Ovaries from progeny flies were dissected and stained with blue precipitate-forming X-Gal in chromogenic β-Galactosidase assays as previously described (Handler et al., 2011). The samples were imaged on an Axio Imager.Z2 widefield microscope with an Axiocam 506 color camera for X-gal stainings.

### RNA experiments

#### poly(A)-enriched RNA-seq of ovaries

Sequencing of polyA RNA-seq libraries of ovaries from *Tj> vreteno^GD^* and *Tj> arrestin2^GD^* flies was performed in (Senti et al., 2023) and analyzed in this manuscript for specific TEs including the MDG1 group. For *flam^Δ412-St2^* RNA-seq, total RNA was extracted with TRIzol (Invitrogen) from ovaries of 3 independent replicates of *flam^Δ412-St2^*;*iso-1/iso-1*;; line #1 and line #2 flies, and 4 independent replicates from control *FRT*;*iso-1/iso-1*;; flies (total 10 samples). RNA was treated with DNase I (ThermoFisher Scientific) and purified using RCC25 kit (Zymo Research) with on-column Zymo DNaseI according to kit instructions. Eluted RNA was polyA-selected twice using magnetic Dynabeads Oligo (dT)_25_ (ThermoFisher Scientific) and subjected to NEBNext Ultra II strand-specific RNA-seq libraries preparation (NEB). Quality control of the RNA and the final libraries was evaluated using Agilent Fragment Analyzer and Qubit fluorometer (ThermoFisher Scientific). Libraries were sequenced using NovaSeqX1B. Reads were mapped, quantified and visualized as described in (Baumgartner et al., 2022).

#### Small RNA-seq libraries of ovaries

Small RNA-seq libraries of two independent replicates of 5 pairs of ovaries from *flam^Δ412-St2^*;;; line #1 and line #2 and control *FRT*;;; flies were generated as previously described (Grentzinger et al., 2020), by isolating Argonaute-sRNA complexes using TraPR ion exchange spin columns. 3′ adaptors containing six random nucleotides plus a 5 nt barcode on their 5′ end and 5′ adaptors containing four random nucleotides at their 3′ end were subsequently ligated to the small RNAs before reverse transcription, PCR amplification, and sequencing on an Illumina NovaSeqX1B. Raw reads were trimmed and mapping was done as in (Baumgartner et al., 2022).

#### Long-read direct RNA-sequencing of OSCs and ovaries

OSCs were grown as in (Niki et al., 2006; Saito et al., 2009) and transfected with siRNA targeting PIWI and control siRNA targeting GFP using Cell Line Nucleofector kit V (Amaxa Biosystems) with the program T-029. siRNA sequences are found in Supplementary table 2. After two days the cells were transfected again and harvested two days later. For ovaries of *Tj> vreteno*^GD^ and *Tj>arrestin2*^GD^ flies, approximately 20 flies per sample were dissected, the ovaries washed with PBS, flash-frozen, and stored at −80°C until processing.

RNA was extracted by homogenizing the samples in TRIzol, and performing phase separation with chloroform, followed by centrifugation at 4816 g for 10 minutes. RNA was precipitated with isopropanol, centrifuged at 20,000 g for 1 hour at 4°C, washed with 75% ethanol, air-dried, and dissolved in nuclease-free water. Poly(A) RNA was enriched using oligo(dT) Dynabeads, with RNA incubated at 80°C, snap-cooled, and bound to beads in binding buffer. After washing, poly(A) RNA was eluted in water at 80°C and quantified using Qubit RNA BR Assay. Libraries were prepared using the ONT RNA002 Direct RNA Library Prep Kit (SQK-002) following the manufacturer’s instructions. Briefly, the ONT RNA adapter was ligated using the NEBNext Quick Ligation Kit, followed by reverse transcription with SuperScript III. After purification with Ampure beads, a second ligation step added the ONT sequencing adapter. Following a final cleanup, libraries were quantified using the Qubit DNA HS Assay, and ∼400 ng of library was used for sequencing. Sequencing was performed on ONT R9 flow cells using MinION Mk1b sequencer. Basecalling was conducted with Dorado v0.8+98ff765, with polyA-tail detection enabled. For TE 3′-end detection, reads with a detected polyA tail were aligned in splice mode using minimap2 to the TE consensus sequences, truncated to their 3′ ends, and analyzed with bedtools genomecov to generate histograms. Visualization of long reads data was done with Tablet (Milne et al., 2013).

#### smFISH of ovaries

smFISH probes targeting the *gag* region in *412, gag-pol* in *Mdg1* and *sORF2-gag-pol* in *Stalker2* sequences were ordered as Stellaris probes for *412* and *Mdg1* (Biosearch Technologies) or as DNA oligos to be labeled in-house (*Stalker2*, IDT) (Gaspar et al., 2017). Probe sequences are listed in Supplementary table 2. smFISH was performed as in (Baumgartner et al., 2022). Briefly, ovaries were dissected, washed and fixed in 4% paraformaldehyde in PBX (PBS with 0.3% Triton-X) at room temperature for 20 minutes. The ovaries were washed and permeabilized with PBX, equilibrated with FISH wash buffer (10% formamide in 2x SSC) and incubated overnight in 50 μL FISH hybridization buffer (100 mg/mL dextran sulfate and 10% formamide in 2x SSC) with 0.5 μL FISH probe at 37 °C. The ovaries were then washed 3x in wash buffer and counterstained with DAPI in 2xSSC, washed with 2XSSC and mounted for imaging.

#### Imaging

All samples for imaging were mounted on slides using ProLong Diamond mounting medium (Invitrogen) and imaged as confocal Z-stacks using Olympus IX3 Series (IX83) inverted microscope, equipped with a Yokogawa W1 spinning disk (SD) of 50 µm pinhole size. Olympus lenses UPLSAPO2 10x/0.4, UPLFLN 40x /0.75, and UPLSAPO 100x /1.4 were used for the acquisition. Diode lasers (OBIS LX, Coherent Corp.) of 405, 488, 561, and 640 nm were used at 100% power (less than 1440, 7970, 7200, and 5700 mW respectively) with the following filters for the emission: 447/60, 525/50, 617/73, 685/40nm. Camera exposure times were set to 300 ms for all channels, and images were acquired with a Hamamatsu ORCA-Fusion (C14440-20UP CMOS) camera. Z-sampling was determined by the optimal Nyquist option in OLYMPUS cellSens Dimension 2.3 software, corresponding to 250 nm in all 100x Z-stacks (65 nm/pixel) and 500 nm in all 40x z-stacks (162.5 nm/pixel). Representative images were then deconvolved using Huygens Professional 24 software (SVI). Deconvolution parameters were individually selected, following the recommendations within the software, with SNR values ranging between 10-30, background subtraction 100 and 30 iterations used. Images were visualized with IMARIS 10 software (Oxford Instruments), setting identical color adjustments to experimental sample images and their respective controls, and individual Z-slices were exported as high-quality TIFF images for the manuscript preparation.

### Protein experiments

#### Generation of polyclonal antibodies

The rat polyclonal antibody targeting 412 Gag (raised against the antigen MLKNDDKTNNAADSE) in co-localization experiments was a kind gift from E. Brasset. Newly generated polyclonal antibodies targeting 412 Gag, 412 sORF1 and 412 sORF2 were raised in rabbits as follows: For Gag, the same antigen was synthesized in-house with N-terminal cysteine residue and coupled to KLH. For sORF1, recombinant His-tagged sORF1 protein was expressed in *E. coli* BL21(DE3), purified by Ni-NTA affinity chromatography followed by ion-exchange chromatography. For sORF2, the antigen from the extracellular region (α-sORF2-N, DTKTVDSTANVINKVEVQNAD) and the antigen from the cytoplasmic region (α-sORF2-C, LKKRYRSTANGIDNI) were synthesized in-house with terminal cysteine residue and coupled to KLH. The synthetic peptides and the sORF1 protein were injected into rabbits to generate polyclonal antisera using the Speedy 28 days protocol (Eurogentec) and purified using glycine elutions of the peptide synthesis columns (Gag, sORF2) or of bound recombinant protein (sORF1).

#### Whole-mount immunofluorescence of ovaries

Immunofluorescence of dissected ovaries was done as in (Baumgartner et al., 2022), and the primary antibodies were used in overnight incubations at 4 °C in the following concentrations: rat anti-Gag (1:1,000), rabbit anti-Gag (1:500), α-sORF1 (1:500), α-sORF2-N (1:250), α-sORF2-C (1:500), rat monoclonal anti-HA (ROAHAHA, Roche) 1:1,000. For labeling F-actin, Alexa Fluor® 488 phalloidin (Invitrogen) was used together with the secondary antibodies at 1:500 and incubated overnight at 4 °C. For labeling DNA, DAPI was used at 1:10,000 for 10 minutes at room temperature.

### Mass-spectrometry

#### Sample preparation for mass-spectrometry experiments

Dissected ovaries from somatic piRNA pathway knockdown (*Tj>vreteno*^GD^) and control knockdown *(Tj>arrestin2*^GD^) flies were dissolved in 500 µL methanol and then protein was precipitated by adding 1mL chloroform. After centrifugation (10 min 10,000 rcf) the supernatant was removed. The pellet was lysed in 10M urea 50mM HCl and then 1M TEAB was added to a final concentration of 100mM. For the reduction of the cysteines DTT and for the alkylation of the free cysteines IAA was used. The samples were diluted with 100mM TEAB Buffer to a urea concentration of 6M and digested with LysC with an enzyme-to-protein ration of 1:50 for 3h at 37 °C. After the pre-digestion with LysC (Wako) the sample was diluted with 100mM TEAB buffer to final urea concentration of 2M. The tryptic digest (Trypsin Gold, Promega) was done with an enzyme-to-protein ration of 1:50 for overnight at 37°C. After the pre-digestion with LysC (Wako) the sample was diluted with 100mM TEAB Buffer to final urea concentration of 2M. The tryptic digest (Trypsin Gold, Promega) was done with an enzyme-to-protein ration of 1:50 for overnight at 37°C.

For OSCs, somatic PIWI knockdown was done with siRNAs as for long-read direct sequencing, using siRNA against GFP as a control. For the digest of the sampled the iST 96x Kit (PreOmics GmbH) was used and performed according to the manufacturer’s protocol.

#### NanoLC-MS Analysis

The nano-HPLC system used was an UltiMate 3000 RSLC nano system (Thermo Fisher Scientific, Amsterdam, Netherlands) coupled to a Q Exactive HF-X or an Orbitrap Eclipse Tribrid mass spectrometer (both Thermo Fisher Scientific, Bremen, Germany), equipped with a Proxeon nanospray source (Thermo Fisher Scientific, Odense, Denmark). Peptides were loaded onto a trap column (Thermo Fisher Scientific, Amsterdam, Netherlands, PepMap C18, 5 mm × 300 μm ID, 5 μm particles, 100 Å pore size) at a flow rate of 25 μL min-1 using 0.1% TFA as mobile phase. After 10 min, the trap column was switched in line with the analytical column (Thermo Fisher Scientific, Amsterdam, Netherlands, PepMap C18, 500 mm × 75 μm ID, 2 μm, 100 Å). Peptides were eluted using a flow rate of 230 nl min-1, and a binary 2h or 3h gradient respectively, which results in a total run time of 140 min or 225 min respectively.

The gradient starts with the mobile phases: 98% A (water/formic acid, 99.9/0.1, v/v) and 2% B (water/acetonitrile/formic acid, 19.92/80/0.08, v/v/v), increases to 35% B over the next 120 or 180 min, followed by a gradient in 5 min to 90%B, stays there for 5 min and decreases in 2 min back to the gradient 98%A and 2%B for equilibration at 30 °C.

The Q Exactive HF-X mass spectrometer was operated in data-dependent mode, using a full scan (m/z range 380-1500, nominal resolution of 60,000, target value 1E6) followed by 10 MS/MS scans of the 10 most abundant ions. MS/MS spectra were acquired using normalized collision energy of 28, isolation width of 1.0 m/z, resolution of 30.000 and the target value was set to 1E5. Precursor ions selected for fragmentation (exclude charge states unassigned, 1, 7, 8, >8) were put on a dynamic exclusion list for 60 s. Additionally, the minimum AGC target was set to 5E3 and intensity threshold was calculated to be 4.8E4. The peptide match feature was set to preferred and the exclude isotopes feature was enabled.

The Eclipse was operated in data-dependent mode, performing a full scan (m/z range 375-1500, resolution 120,000, target value 1E6, normalized AGC target 250%) at 3 different compensation voltages (CV-45, - 55, −75), followed each by MS/MS scans of the most abundant ions for a cycle time of 1 sec per CV. MS/MS spectra were acquired using an isolation width of 1 m/z, AGC target value of 30,000, normalized AGC target 300%, minimum intensity of 50,000, maximum injection time 15 ms, activation type.

HCD with a collision energy of 30 %, using the Iontrap for detection in the scan mode Turbo. Precursor ions selected for fragmentation (included charge states 2-6) were excluded for 40 s. The monoisotopic precursor selection filter and exclude isotopes feature were enabled.

#### Mass-spectrometry data processing

For peptide identification, the RAW-files were loaded into Proteome Discoverer (version 2.5.0.400, Thermo Scientific). All hereby created MS/MS spectra were searched using MSAmanda v2.0.0.16129, Engine version v2.0.0.16129 (Dorfer et al., 2014). The RAW-files were searched against the *Drosophila melanogaster* database of Flybase, version r6.36 (22,226 sequences; 20,310,919 residues) and a TE database, which includes all BDGP consensus sequences (Kaminker et al., 2002) translated in six reading frames with a minimum open reading frame length of 40 amino acids, supplemented with common contaminants, using the following search parameters: Iodoacetamide derivative on cysteine was set as a fixed modification, oxidation on methionine and deamidation on asparagine and glutamine were set as variable modifications. The peptide mass tolerance was set to ±5 ppm and the fragment mass tolerance to ±8 ppm for the QEx-HFX-measurement, as well as ±5 ppm and fragment mass tolerance of ±500 mmu for the Eclipse-measurement. The maximal number of missed cleavages was set to 2, using tryptic enzymatic specificity. The result was filtered to 1 % FDR on protein level using Percolator algorithm (Käll et al., 2007) integrated in Thermo Proteome Discoverer.

The localization of the post-translational modification sites within the peptides was performed with the tool ptmRS, based on the tool phosphoRS (Taus et al., 2011).

Protein areas have been computed in IMP-apQuant (Doblmann et al., 2019) by summing up unique and razor peptides. Resulting protein areas were normalized using iBAQ (Schwanhäusser et al., 2011) and sum normalization was applied for normalization between samples.

### Electron microscopy and tomography

#### Preparation for ultrastructural preservation and tomography

*Drosophila flam^Δ412-St2^*;*iso-1* and *FRT;iso-1* ovaries were dissected in ice-cold PBS and mounted in a 200µm recess of an aluminum carrier with a diameter of 3mm, covered with the flat side of a carrier, and immediately high pressure frozen in a HPF Compact 01 (Engineering Office M. Wohlwend GmbH, Switzerland). To avoid air bubbles, 5% BSA in Sörensen phosphate buffer was used as filling material. Subsequently the samples were processed in a Leica AFS-2 automated freeze substitution unit (Leica Microsystems, Austria), following this protocol: 60h at –90°C, warm up at a rate of 2°C per hour to –54°C, 18h at –54°C, warm up at a rate of 5°C per hour to –24, 15h at –24°C, warm up at a rate of 6°C per hour to 20°C, 5h at 20°C. Over this 5d-period, the frozen water in the samples was substituted with acetone containing 1% osmium tetroxide, 0.2% uranyl acetate and 5% water (UA from Merck, Germany; OsO4 from EMS, USA). The dehydrated samples were washed 3 times in anhydrous acetone at RT. Samples were infiltrated with a medium hard mixture of Epoxy resin (Agar 100 from Agar Scientific, UK), in a graded series of acetone and resin (2:1, 1:1, 1:2 for 1h each), followed by pure resin for a few hours and polymerization in the oven for 2d at 60°C.

#### TEM ultrastructure imaging of epon-fixed samples

The obtained resin blocks were cut using a Leica UCT ultramicrotome (Leica Microsystems, Austria) at a nominal thickness of 50-70nm and the sections were picked up on 100mesh copper grids or slot-grids (Agar Scientific, UK), previously coated with a self-made Formvar® support film (Formvar powder from Agar Scientific, UK). Sections obtained from high pressure freezing/freeze substitution, did not need any further contrasting. Grids were then examined in a FEI Morgagni 268D transmission electron microscope, operated at a high tension of 80kV (FEI/Thermo Fisher, The Netherlands). Digital images were acquired using a Megaview III CCD camera (Olympus-SIS, Germany). For quantification of capsid diameter, 120 capsids from different images taken at different time points were measured manually and the average diameter represents the most frequently observed capsid size.

#### Electron Tomography

Epon-fixed samples were sectioned into 250 nm sections, and a concentrated solution of 10 nm gold beads (Aurion, The Netherlands) was applied in a single drop and incubated for a period of three minutes. Double-axis tilt series were imaged using a Tecnai G2 20 microscope equipped with an Eagle 4 k HS CCD camera (FEI, the Netherlands). The microscope was operated at 200 kV and the SerialEM software (Mastronarde, 2005) was employed for data acquisition. To prevent shrinkage during the acquisition of tilt series data, the section was subjected to low-magnification cooking for a period of 20 minutes prior to data acquisition. Dual axis tilt series were recorded at 1° increments with a tilt range of −57°/+52° and −60°/+59°, respectively. The raw data were processed into tomograms using the IMOD software (Mastronarde & Held, 2017).

#### Cryo-immunoEM - Tokuyasu method

*Drosophila flam^Δ412-St2^*;*iso-1* and *FRT;iso-1* ovaries were dissected in ice-cold PBS and fixed in 2% paraformaldehyde and 0.2% glutaraldehyde (both EM-grade, EMS, USA) in 0.1 M PHEM buffer (pH 7) for 2h at RT, then overnight at 4°C. The fixed ovaries were embedded in 12% gelatin and cut into 1 mm^3^ blocks which were immersed in 2.3 M sucrose overnight at 4°C. These blocks were mounted onto Leica specimen carrier (Leica Microsystems, Austria) and frozen in liquid nitrogen. With a Leica UCT/FCS cryo-ultramicrotome (Leica Microsystems, Austria) the frozen blocks were cut into ultra-thin sections at a nominal thickness of 70nm at −110°C. A mixture of 2% methylcellulose (25 centipoises) and 2.3 M sucrose in a ratio of 1:1 was used as a pick-up solution. Sections were picked up onto 200 mesh Ni grids (Gilder Grids, UK) with a carbon coated formvar film (Agar Scientific, UK). Fixation, embedding and cryo-sectioning as described by (Tokuyasu, 1973). Prior to immunolabeling, grids were placed on plates with solidified 2% gelatine and warmed up to 37 °C for 20 min to remove the pick-up solution. After quenching of free aldehyde-groups with glycine (0.1% for 15 min), a blocking step with 1% BSA (fraction V) in 0.1 M Sörensen phosphate buffer (pH 7.4) was performed for 40 min. The grids were incubated in primary antibody (α-Gag and α-sORF2-C), diluted 1:50 in 0.1 M Sörensen phosphate buffer over night at 4°C, followed by a 2h incubation in the secondary antibody, a goat-anti-rabbit antibody coupled with 6 nm gold particles (GAR 6 nm, Aurion, The Netherlands), diluted 1:20 in 0.1 M Sörensen phosphate buffer, performed at RT. The sections were stained with 4% uranyl acetate (Merck, Germany) and 2% methylcellulose in a ratio of 1:9 (on ice). All labeling steps were done in a wet chamber. The sections were inspected using a FEI Morgagni 268D TEM (FEI/Thermo Fisher, The Netherlands) operated at 80kV. Electron micrographs were acquired using a Megaview III CCD camera (Olympus-SIS, Germany).

### Computational analysis

#### Annotation and phylogenetic analysis of LTR retrotransposon sequences

Consensus sequences and annotated features (LTRs, *gag* and *pol* ORFs, sORFs) for all *D. melanogaster* LTR retrotransposons studied in this work were collected from the BDGP v.9.4.1 database (Kaminker et al., 2002), Repbase (Bao et al., 2015) and primary literature (Avedisov et al., 1990; Cherkassova et al., 1991; Costas et al., 2001; Makarova, 1997; Mugnier et al., 2005; Yuki et al., 1986). *sORF1* and *sORF2* were identified as predicted 60-130 amino acids long ORFs, positioned between the 5’ LTR and *gag* and in the same orientation as *gag*. The absence of intact *env* ORF was determined by measuring the distance between the end of *pol* and the start of the 3’ LTR for the MDG1 group retrotransposons. The phylogenetic tree shown in Fig. 1A was generated by aligning the *pol* sequences from the protease domain to and including the conserved integrase domain as in (Senti et al., 2023), and visualized with iTol v6 (Letunic & Bork, 2021). For *Blood* the consensus sequence in Repbase and BDGP shows only the possible translation of the first 11 amino acids of sORF2, however the sORF2 of *Blood* was determined based on analysis of five heterochromatic *Blood* insertions which show an intact sORF2, but have a defective *gag*, *pol* or interrupted sequence by “nested” TEs.

#### *Flamenco* enrichment of TE groups analysis

Non-overlapping genomic windows of 100 kb were generated from the *Drosophila melanogaster* reference genome (dm6), excluding mitochondrial and Y chromosome sequences. To precisely capture the *flamenco* locus, a TE-rich region on the X chromosome (*chrX: 21,631,891–21,931,891*), windows were centered to align exactly with this region. TE content within each window was quantified using RepeatMasker annotations (Smit, A.F.A. Hubley, R., & Green, P., 2013), irrespective of strand orientation. Windows with >50% TE content were classified as heterochromatic tiles and retained for further analysis. TE content was further stratified by strand and TE category (e.g., LTR, LINE, DNA) and visualized in Fig. 1C. Statistical significance for the enrichment of specific TE groups, such as the MDG1 group, was assessed by calculating a z-score and an empirical p-value (*e.g.*, MDG1: *z* = 24, p = 1.45 × 10-2).

#### Alphafold3 predictions

AlphaFold3 (Abramson et al., 2024) was used to predict protein structures of sORF1 and sORF2 and structures were analyzed using UCSF ChimeraX (Pettersen et al., 2021).

#### Multiple sequence alignment and sORF2/FAST-like structural features prediction

Amino acid sequences of proteins were aligned using MAFFT (Madeira et al., 2024) and visualized with Jalview (Waterhouse et al., 2009). Manual adjustments were introduced to clarify demonstrated or presumed structural and functional motifs characteristic of FAST proteins. Color scheme is according to ClustalX conservation criteria. The transmembrane domain and extracellular/cytoplasmic topology were predicted using TMHMM (Krogh et al., 2001). N-myristoylation prediction was done with GPS-Lipid for the initial identification of sORF2 proteins in *Drosophila* (Xie et al., 2016), and for all other analyses determined by presence of the consensus MGxxxS/T motif. A polybasic region was identified by having 3 or more K or R amino acids within the first 15 amino acids immediately following the transmembrane domain. Amphipathic helices were predicted and hydrophobic moment <µH> calculated using Heliquest (Gautier et al., 2008).

#### Identification of insect sORF2/FAST-like candidates

A database of 8,417 consensus sequences of *Gypsy* LTR retroviruses in arthropods was downloaded from Repbase (March 2024). All interior sequences were translated in all six reading frames and screened for sORF2-like proteins using the following criteria: ORF size between 60-210 amino acids (180-630 nt); a single transmembrane domain predicted by TMHMM 2.0c and Phobius 1.01 (Käll et al., 2004); N-terminal extracellular (NONCYTO) and C-terminal intracellular (CYTOPLASMIC) topology; an ectodomain containing an N-myristoylation motif (MGxxxS/T) with a length between 10-50 amino acids; and a cytoplasmic region smaller than 150 amino acids. This screening identified 105 candidate sequences, which were further analyzed for their genomic coordinates within the Repbase consensus sequence (fig. S10) and the taxonomic classification of the arthropod genomes, visualized with iTol (Figure 5B). Alignments and feature annotations (e.g. polybasic, amphipathic motifs) were performed as described above.

#### Genome-wide scanning of TE insertions in select genomes

To identify individual TE insertions, RepeatMasker version 4.1.7-p1 (Smit, A.F.A. Hubley, R., & Green, P., 2013) was used to screen LTR retrotransposons in the following genomes with custom TE consensus sequence libraries: *D. melanogaster* dm6 (GCF_000001215.4, Dfam database (Storer et al., 2021) using RepeatMasker default parameters-species 7227 rmblastn version 2.14.1 + FamDB: CONS-Dfam_3.8), *A. aegyptii* L5 (GCF_002204515.2) with TE library from (Daron et al., 2024). For *S. oryzae* 2.0 (GCF_002938485.1) TE annotations were taken from (Parisot et al., 2021). Genomic nucleotide sequences were screened for sORF2/FAST-like candidates as described for the Repbase consensus sequences. Resulting candidates were filtered for minimal overlap 20 nt with any RepeatMasker predicted repetitive regions. To match the individual insertions to the identified sORF2/FAST-like candidates from Repbase consensus sequences, BLAST searches on the protein sequences were performed with default parameters, and results were filtered by an E-value cutoff of 1e-10.

#### Identification of sORF2/FAST like proteins in non-enveloped viruses

The 105 sORF2/FAST-like amino acid sequences identified in insect LTR retrotransposons were aligned using PRANK (Löytynoja, 2014) and an HMM profile was constructed using HMMER v3.4 (Eddy, 2011). This HMM profile was used to search the NCBI nr database (January 2025) for viral sequences, with human and non-human viruses analyzed separately. For non-human viruses, 56 hits passed the E-value threshold of 0.05, including 39 sequences from picornaviruses and 2 from nackednaviruses. All picornavirus genomes identified with this search that were between 1,500 and 20,000 nt in length are shown in Supplementary Figure S11. Select sequences were aligned and shown in Figure 5E, including Goose megrivirus A (NC_033793.1), Duck megrivirus A2 (KC663628.1), Swan megrivirus A, Chicken megrivirus C (KF961187.1), Turkey melegrivirus A (KF961188.1), Peafowl picornavirus sp. (MT138340.1), and Chickadee poecivirus A1 (KU977108.1).

For nackednaviruses, all 13 genome sequences lacking *preS/S* gene were extracted from Supplementary file S1 of (Lauber et al., 2017) and shown as genome annotations in Supplementary Figure S11. sORF2/FAST-like features were predicted using the same criteria as for insect LTR retrotransposon candidates. Select sequences were aligned and shown in Fig. 5F (African cichlid ACNDV, European eel EENDV, Rockfish RNDV, Astatotilapia ANDV, Rainwater killifish KNDV-Lp-1).

---

**Supplementary Figure 1:**
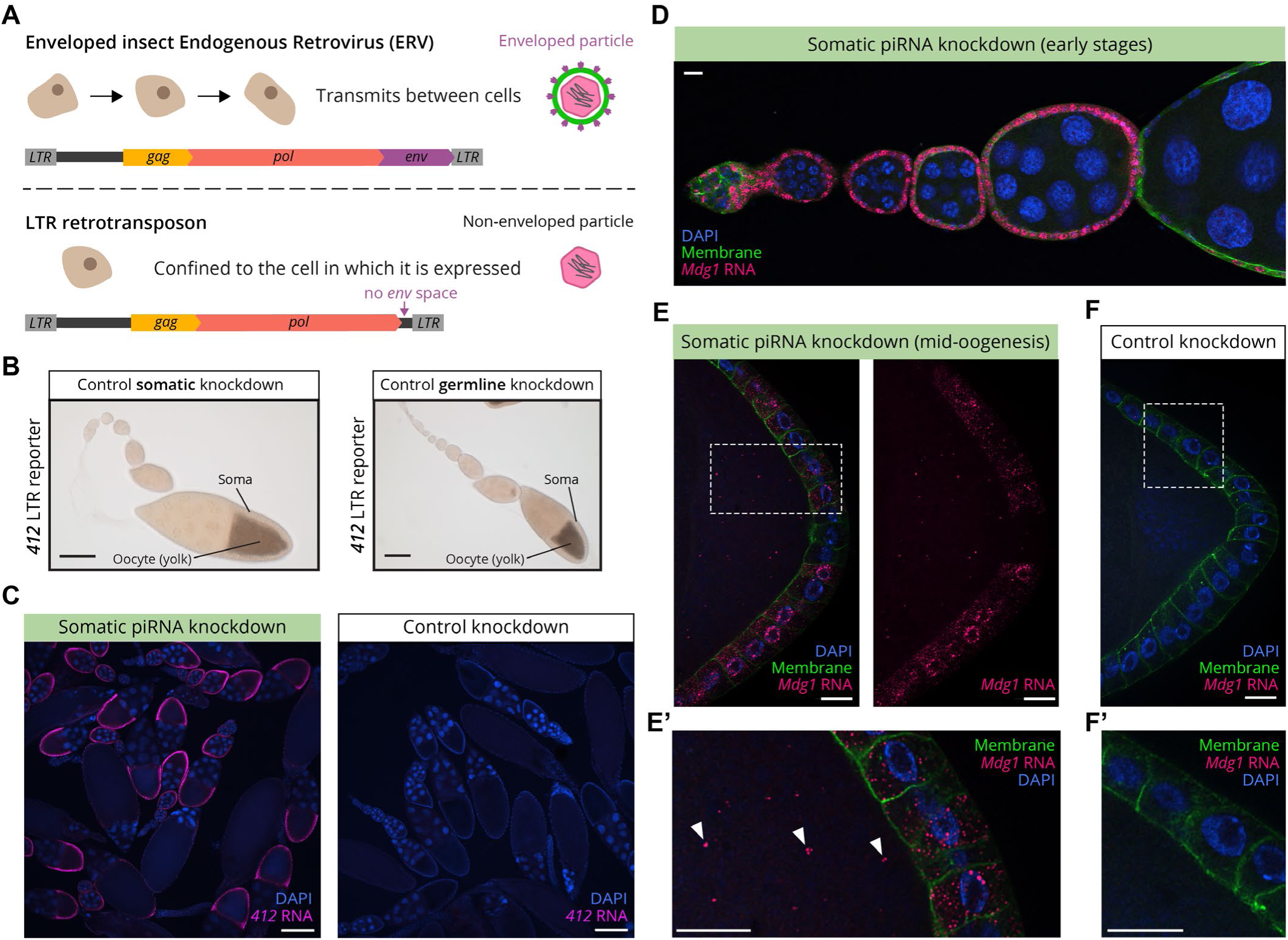
Robust somatic expression and oocyte transmission of MDG1 retrotransposons. **(A)** Schematic comparison of insect ERV and LTR retrotransposon genome organization. Insect ERVs encode an Envelope protein (purple) that enables cell-to-cell transmission, while LTR retrotransposons lack an *env* gene and are typically confined to the cell in which they are expressed. **(B)** Related to Figure 1E - control knockdowns of the transcriptional *412 LTR-lacZ* reporter, left: somatic piRNA pathway knockdown (*Tj>Gal4,arr2-RNAi*), right: germline piRNA pathway knockdown (*MTD>Gal4,wsh-RNAi*). Scale bar: 100 µm. **(C)** Left: Overview of somatic piRNA pathway knockdown (*Tj>Gal4,vret-RNAi*) sample showing *412* smFISH (magenta) signal in somatic follicle cells across multiple ovarioles. Right: overview of *412* smFISH signal in control knockdown (*Tj>Gal4,arr2-RNAi*) ovarioles. **(D-F)** smFISH staining of **(D)** early and **(E)** mid-oogenesis follicles for LTR retrotransposon *Mdg1* (pink), recapitulating the soma-to-oocyte transfer seen for additional MDG1 group retrotransposons in the somatic piRNA pathway knockdown background (*Tj>Gal4,vret-RNAi*, left) whereas in **(F)** control ovaries *Mdg1* is not expressed (*Tj>Gal4,arr2-RNAi*). White arrowheads denote RNA signals within the oocyte. In (D-F) somatic membranes are marked by myristoylated GFP. Scale bars (C-F): 10 µm.

**Supplementary Figure 2:**
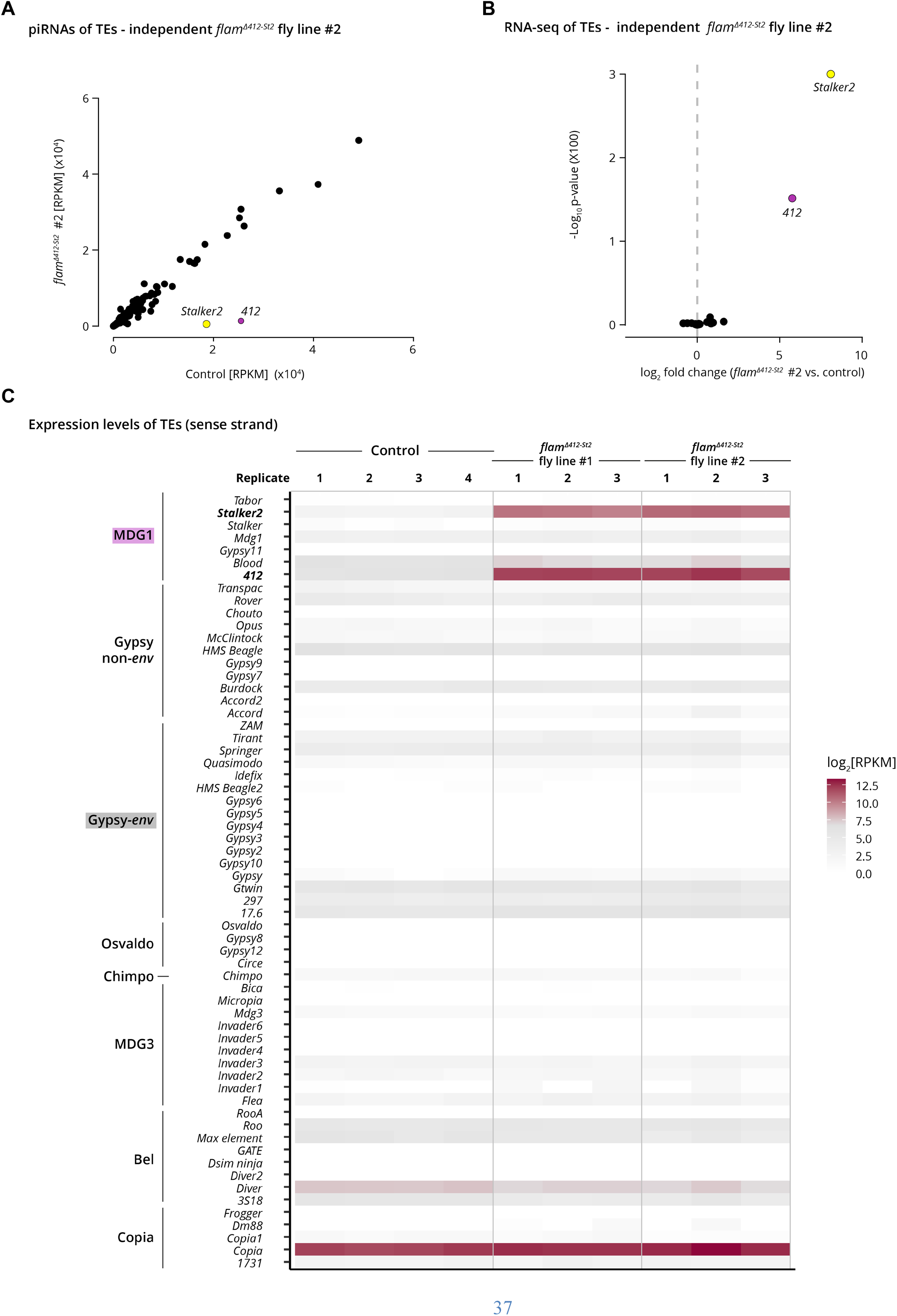
RNA expression of TEs in *flam^Δ412-St2^* ovaries. **(A)** Small RNA sequencing of piRNAs from *flam^Δ412-St^* ovaries (homozygous mutant line #2), compared to control ovaries (genetically identical except for the targeted *flamenco* deletion). Each datapoint represents the number of antisense reads mapped to a TE consensus sequence, normalized as reads per kilobase of TE per million mapped microRNAs (RPKM). **(B)** Volcano plot of whole transcriptome poly(A)-tailed RNA-seq, from *flam^Δ412-St2^* ovaries(homozygous mutant line #2), compared to control ovaries (n=3). Each datapoint represents the number of reads mapped to a TE consensus sequence. **(C)** Heatmap of expression levels [RPKM] for 62 LTR retrotransposons across three replicates of *flam^Δ412-St^* ovaries (two independent mutant lines) and four replicates of control ovaries. Values between 0 and 1 are rounded to 1 and represented as white squares. *Copia*, which does not encode for an Envelope, is known to be constitutively highly expressed in ovaries (Klumpe et al., 2025).

**Supplementary Figure 3:**
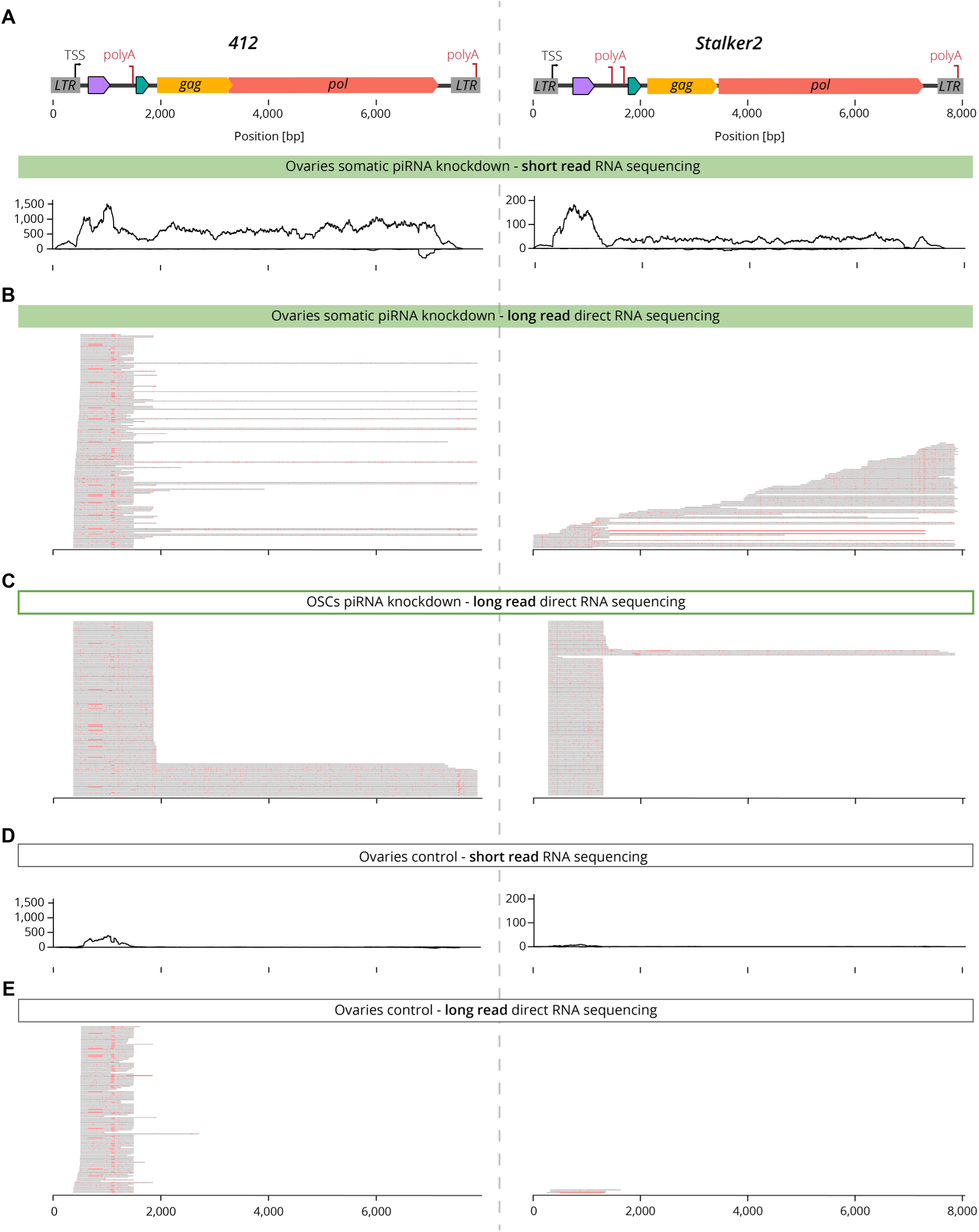
*412* and *Stalker2* produce long and short transcripts. For (A-E), *412* data is shown on the left and *Stalker2* is shown on the right. For (B,C,E) partial subsets of long-read direct RNA sequencing reads are shown, with reads selected to show both long and short isoforms, and sorted according to the first mapped nucleotide of the read. Variant regions deviating from consensus are shown in red. **(A)** Coverage plots of short-reads RNA-sequencing of one representative replicate from somatic piRNA pathway knockdown ovaries (*Tj>Gal4,vret-RNAi*) sample. Y-axis is number of reads mapped to consensus sequences, normalized as reads per kilobase of TE per million mapped reads (RPKM). **(B)** Subset of long-read direct RNA sequencing reads, showing individual reads mapped to the consensus sequence, of the same genotype as in (A). **(C)** Subset of long-read direct RNA sequencing reads from somatic piRNA pathway knockdown in OSCs (treated with siRNA targeting Piwi). **(D)** Coverage plots of short-reads RNA-sequencing of one representative replicate from control knockdown ovaries (*Tj>Gal4,arr2-RNAi*) sample. Y-axisas in (A). **(E)** Subset of long-read direct RNA sequencing reads, showing individual reads mapped to the consensus sequence, of the same genotype as in (D).

**Supplementary Figure 4:**
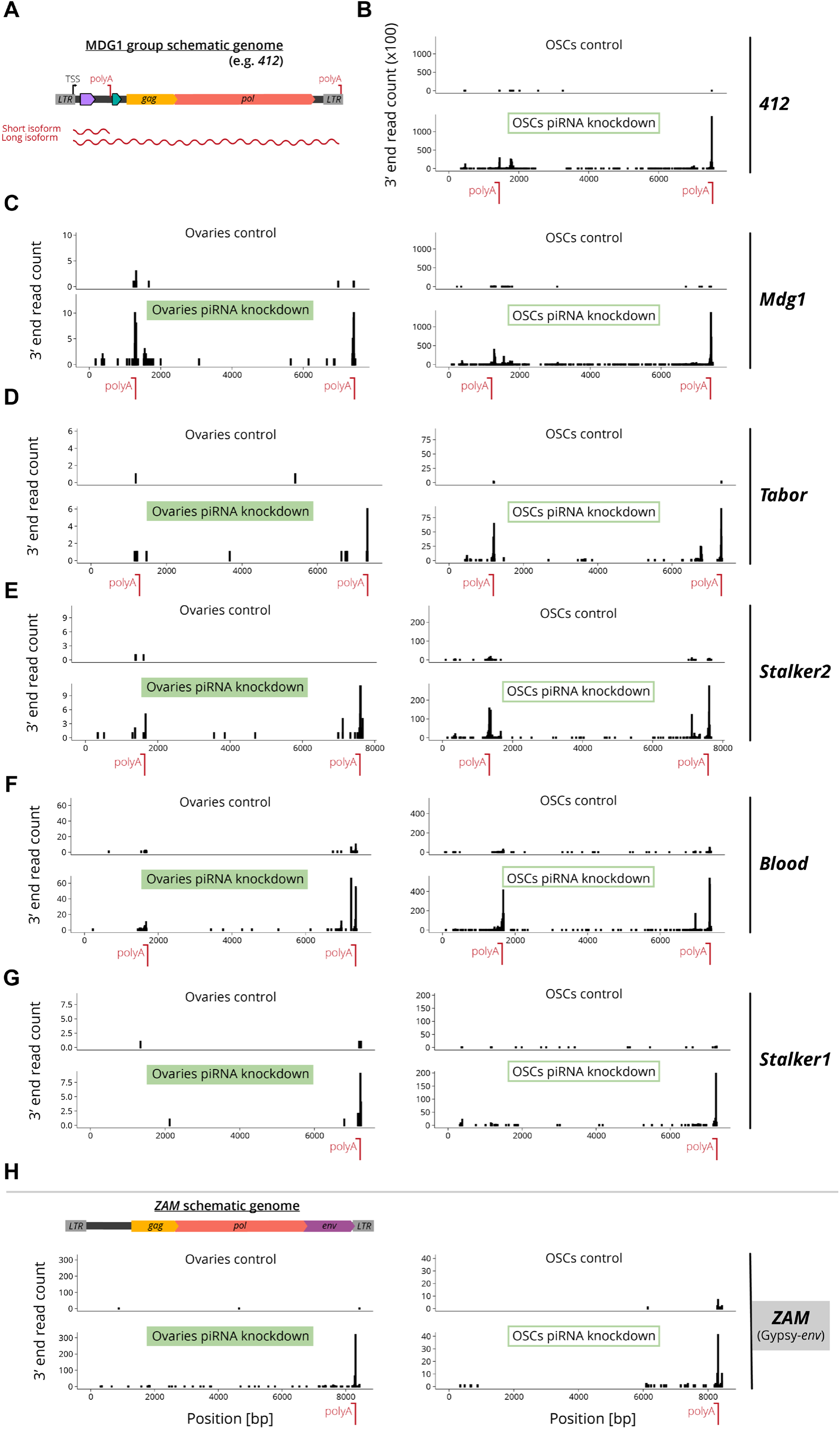
Premature transcription termination sites are found in most MDG1 retrotransposons. **(A)** Schematic representation pf MDG1 LTR retrotransposon genomes, showing ORFs (colored arrows), LTRs (grey boxes), transcription termination sites (“polyA”), and wavy lines representing the short and long isoforms produced based on the polyA site that is predominantly used. **(B-H)** Mapping of 3’ ends of long-read direct RNA-sequencing reads corresponding to the consensus sequences of (B-G) MDG1 retrotransposons, and (H) the enveloped insect-ERV ZAM as a control. Left panels: ovaries, comparing control (*Tj>Gal4, arr2-RNAi*, top) and somatic piRNA pathway knockdown (*Tj>Gal4, vret-RNAi*, bottom). Right panels: OSCs, comparing control (treated with siRNA targeting GFP) and piRNA pathway knockdown (treated with siRNA targeting Piwi). *Stalker1* appears to lack a premature transcription termination site, based on the 3’ end data and sequence analysis. For *412* the ovaries samples are shown in main Figure 3B. The Y-axis indicates the number of reads.

**Supplementary Figure 5:**
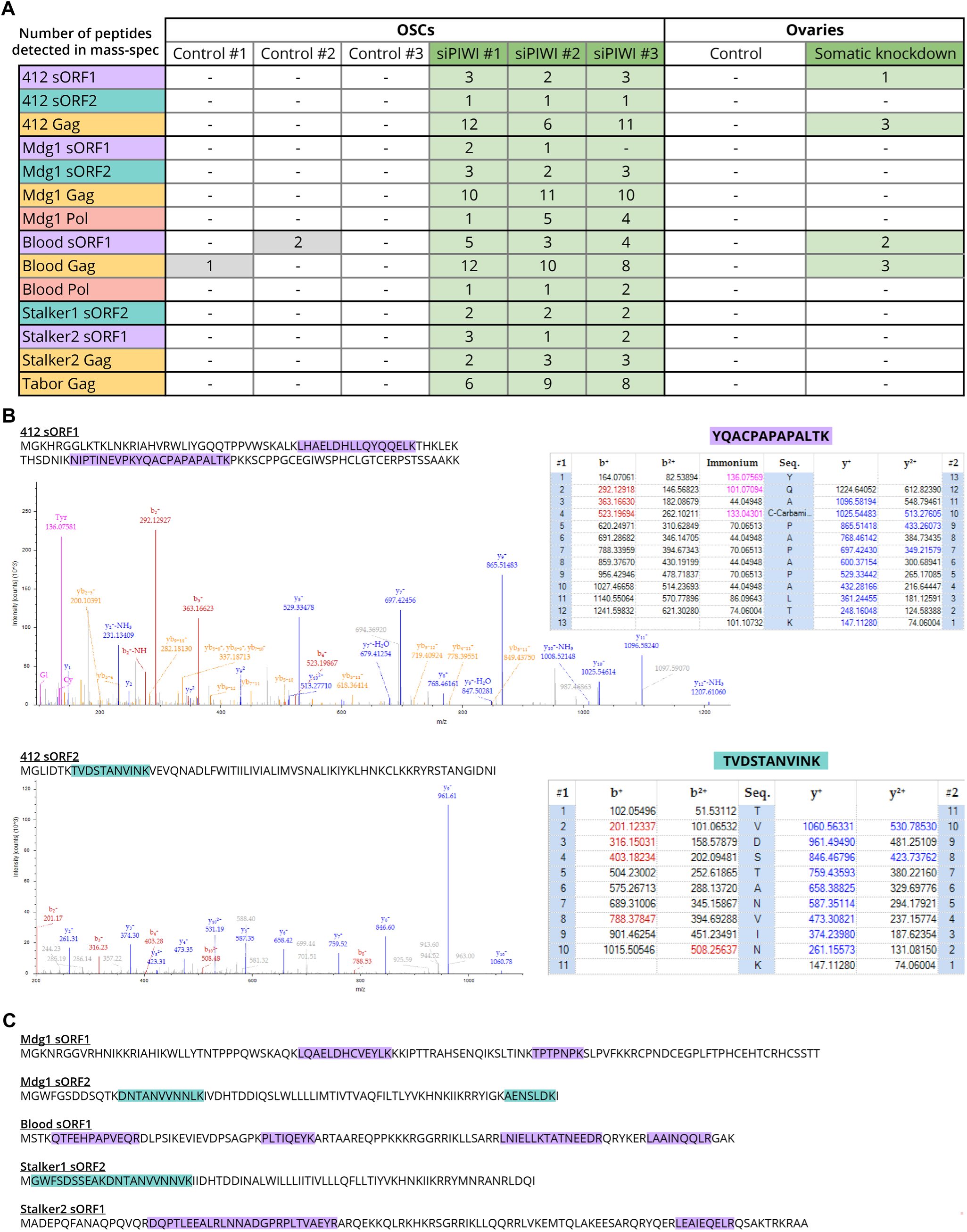
Untargeted proteomics evidence for translation of MDG1 retrotransposon ORFs. **(A)** Mass-spectrometry detection of peptides corresponding to sORF1 (purple), sORF2 (cyan) and Gag (orange) sequences of MDG1 LTR retrotransposons in control OSCs (treated with siRNA targeting GFP), piRNA pathway knockdown OSCs (treated with siRNA targeting Piwi), control ovaries (*Tj>Gal4, arr2-RNAi*) and somatic piRNA pathway knockdown ovaries (*Tj>Gal4, vret-RNAi*). **(B)** Two representative fragment spectra of detected peptides supporting the presence of 412 sORF1 in somatic piRNA pathway knockdown ovaries and sORF2 in piRNA pathway knockdown OSCs. **(C)** Sequences of sORF1 and sORF2 peptides detected in mass-spec, supporting the translation of these proteins from MDG1 retrotransposons.

**Supplementary Figure 6:**
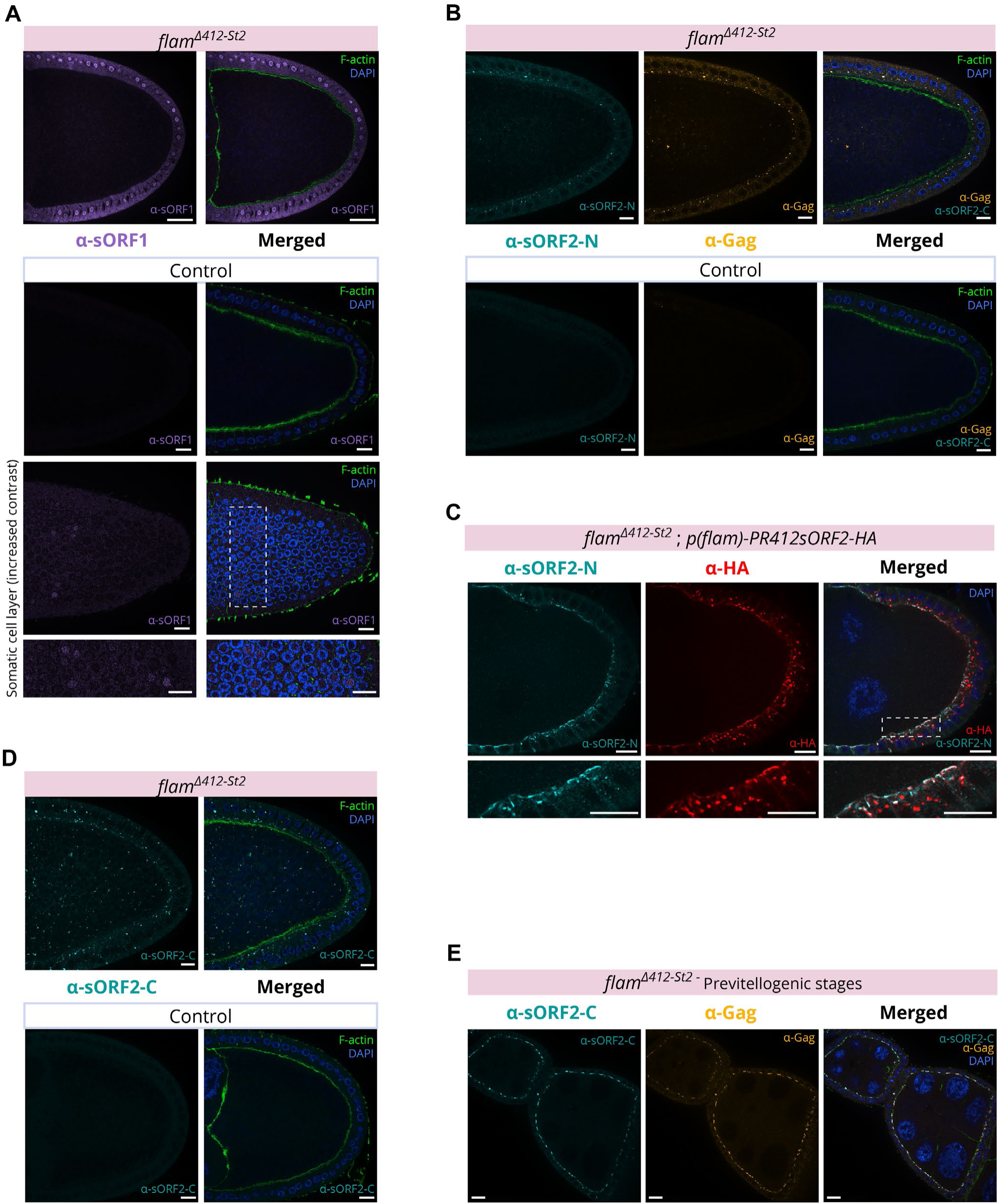
Antibodies targeting sORF1 and sORF2 are specific and support distinct cellular localizations. **(A)** Whole mount immunofluorescence of *flam^Δ412-St2^* ovaries (top) and control ovaries (bottom) with α-sORF1 antibody (purple). A weak nuclear staining of sORF1 is visible in control ovaries with increased contrast settings, consistent with RNA-seq and mass-spec data (bottom panels). **(B)** Whole mount immunofluorescence of *flam^Δ412-St2^* ovaries (top) and control ovaries (bottom) stained with α-sORF2-N (cyan) and α-Gag (orange). **(C)** Whole mount immunofluorescence of *flam^Δ412-St2^* ovaries expressing a C-terminal HA-tagged *412 sORF2* transgene under a *flamenco* promoter, stained with α-sORF2-N (cyan) and α-HA (red), showing good co-localization (white) of the two antibodies on the apical somatic membranes. **(D)** Whole mount immunofluorescence of *flam^Δ412-St2^* ovaries (top) and control ovaries (bottom) with α-sORF2-C (cyan). **(E)** Co-localization of α-sORF2-C (cyan) and α-Gag (orange) on the apical somatic cell membranes in previtellogenic stages. For (A-B, D-E) F-actin is labelled with phalloidin (green) to demarcate cortical actin near the plasma membrane. Scale bars (A-E): 10 µm.

**Supplementary Figure 7:**
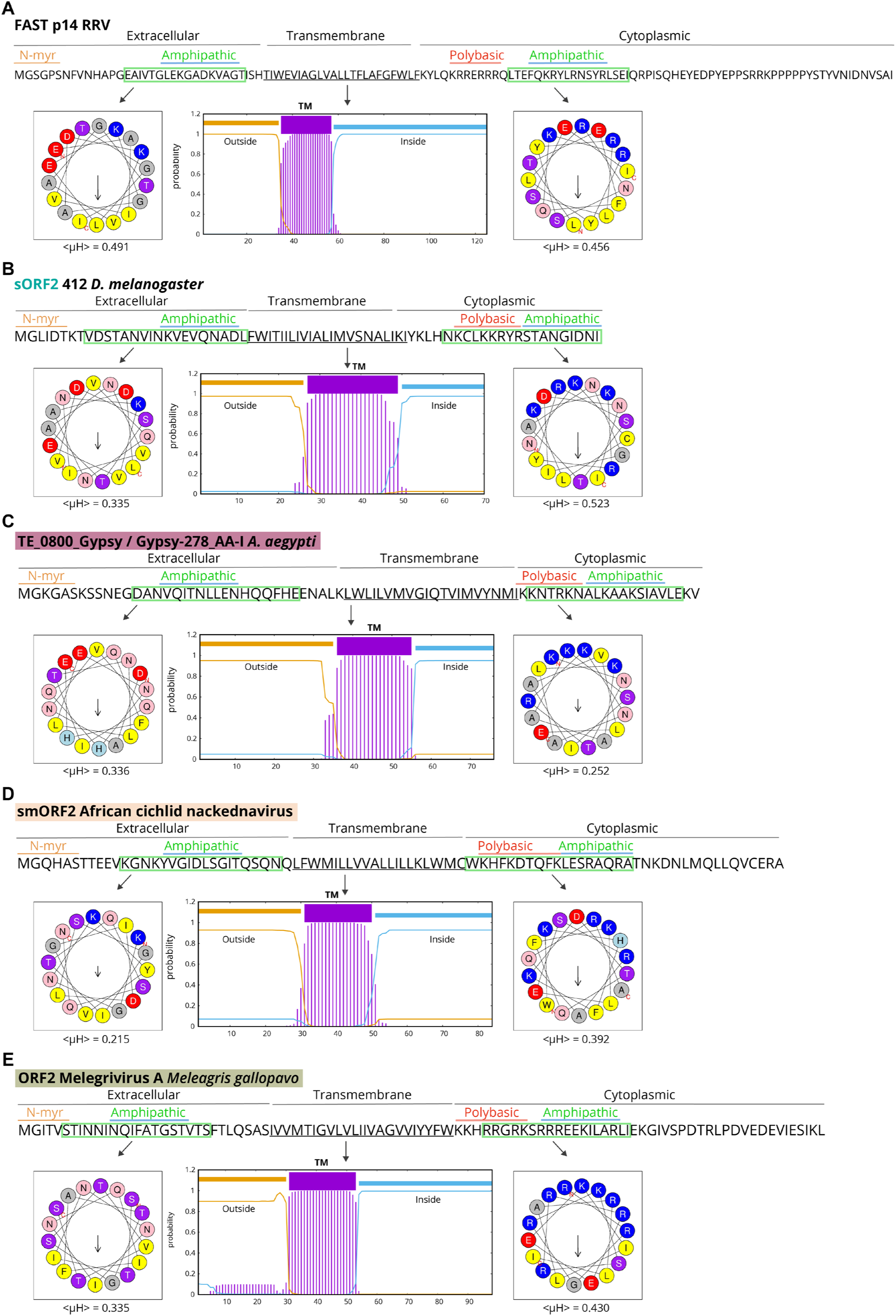
Transmembrane and amphipathic domains in newly discovered fusogen-like proteins. Predictions for amphipathic helices using Heliquest and for transmembrane domains using TMHMM (see Materials & Methods) in (A) known FAST protein p14 from reptilian orthoreovirus (Genbank AAP03134.1), (B) sORF2 from *412* LTR retrotransposon (Genbank X04132), (C) Newly identified ORF within TE_0800_Gypsy/Gypsy-278 from *A. aegyptii* genome (Genbank GCF_002204515.2), (D) smORF2 from African cichlid nackednavirus (Genbank MH158727) and (E) ORF2 from turkey-infecting Melegrivirus A (Genbank KF961188). Helical projections and the hydrophobic moment values <µH> shows various degrees of partitioning of hydrophobic residues (yellow) and polar/charged residues (blue, red, purple, pink) on opposite sides of the helix, corresponding to amphipathic helices.

**Supplementary Figure 8:**
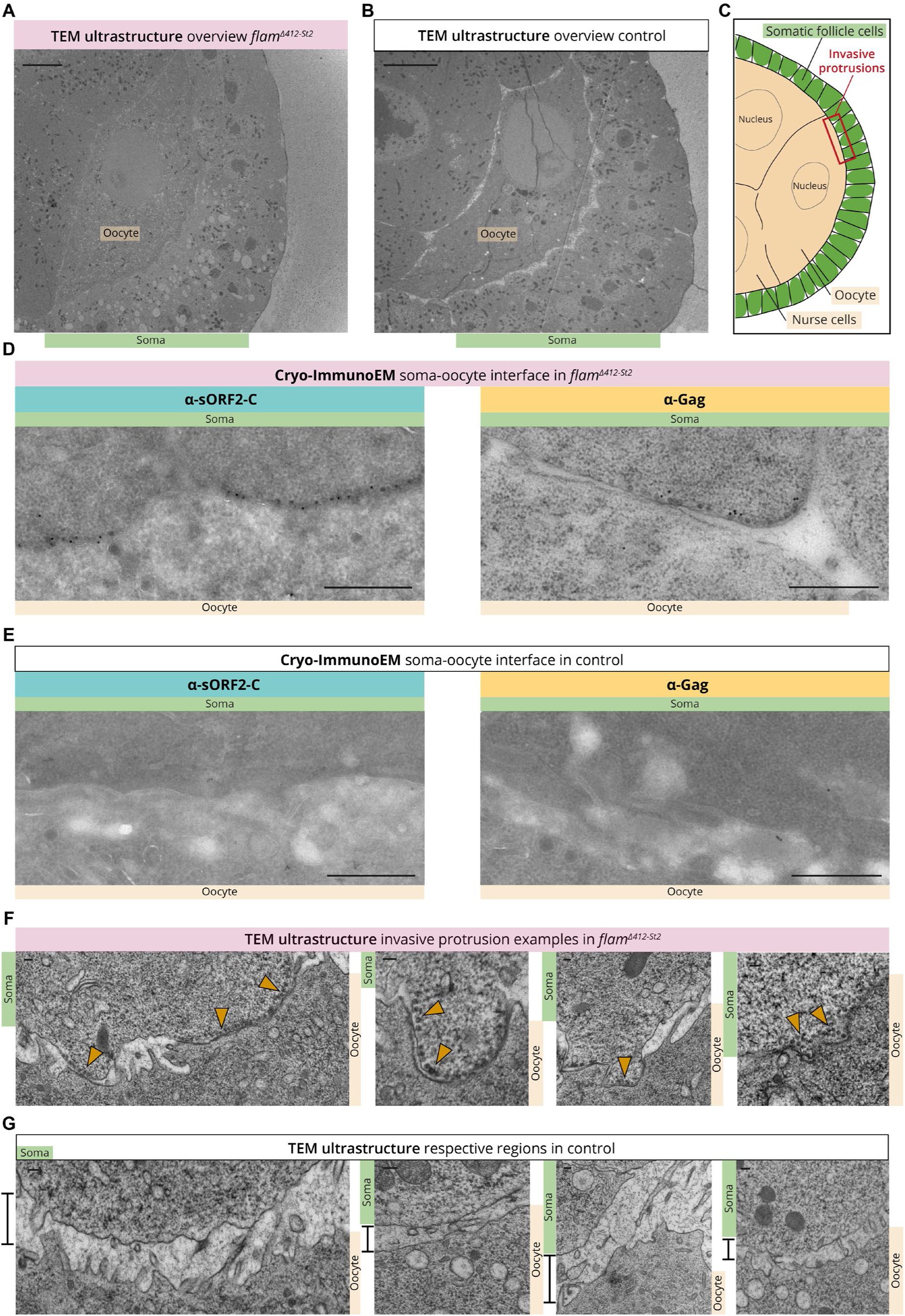
Characterization and validation of observed capsids and protrusions in *flam^Δ412-St2^* follicles. **(A-C)** Overview of region of interest in TEM ultrastructure studies. (A) Stage 7 *flam^Δ412-St2^* follicle does not show any major morphological defects compared to (B) control follicle of the same stage. For (A-B) Scale bar: 5 µm (C) Cartoon of characteristic region (boxed red rectangle) for observed invasive protrusions in the soma-oocyte interface of *flam^Δ412-St2^* follicles. **(D)** Cryo-immunoEM of *flam^Δ412-St2^* stage 7 follicle using α-sORF2-C antibody (left image, gold particles as black dots) and α-Gag (right image, gold particles as black dots), in proximity to accumulated capsids on the apical membranes of somatic follicle cells. **(E)** Same for (D), only for control ovaries. For (D-E) Scale bar: 500 nm. **(F-G)** Observation of soma-oocyte interface in ultrastructure of (F) *flam^Δ412-St2^*follicles and (G) control follicles. Notice the appearance of capsid-filled invasive protrusions (orange arrowheads) which are in close contact with the oocyte membrane in *flam^Δ412-St2^* follicles, as opposed to control follicles lacking such structures and a larger distance between soma and oocyte membranes (black bar). For (F-G) Scale bar: 100 nm.

**Supplementary Figure 9:**
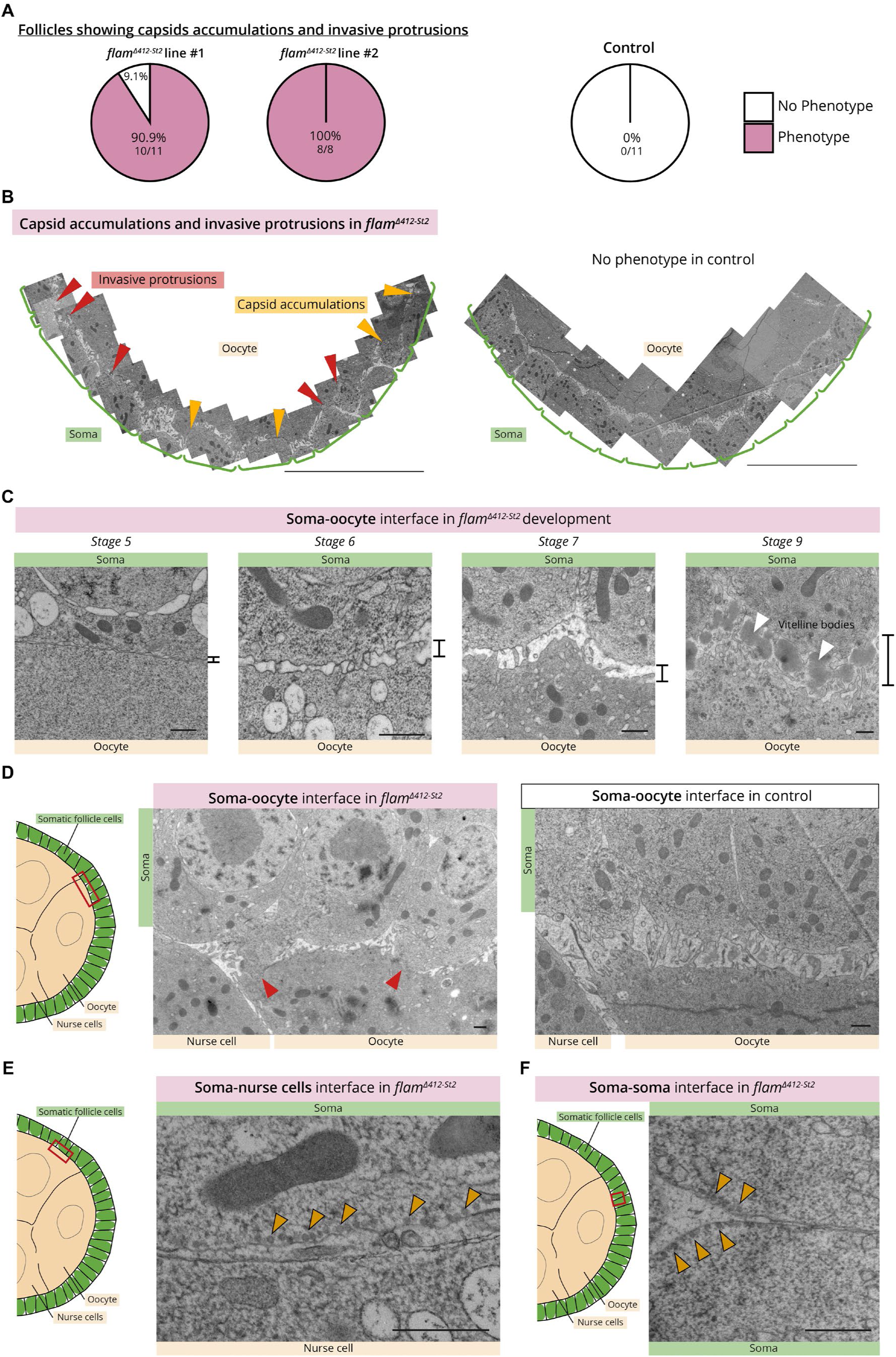
Invasive protrusions form exclusively in the soma-oocyte interface of *flam^Δ412-St2^* follicles during a developmental time window. **(A)** Quantification of follicles exhibiting capsid accumulations and invasive protrusions as observed in TEM ultrastructure of two independent mutant *flam^Δ412-St2^* lines and control ovaries. **(B)** Spatial distribution of capsid accumulations (orange arrowheads) and invasive protrusions (red arrowheads) along the apical membranes of somatic follicle cells (green brackets) facing the oocyte. Images were manually stitched for representative *flam^Δ412-St2^* and control follicles. Scale bar: 10 µm. High-resolution images are available in Supplemental Data files S1 and S2. **(C)** Soma-oocyte interface of *flam^Δ412-St2^* follicles at different developmental stages. Notice in stage 5 the absence of microvilli and close association of somatic and oocyte membranes. As development progresses, somatic and oocyte membranes separate but maintain somatic microvilli across the perivitelline space, until deposited vitelline bodies coalesce to form the protective layer of the egg. **(D)** Invasive protrusions (red arrows) in the soma-oocyte interface of *flam^Δ412-St2^* stage 7 follicle (left), and the respective area in a control follicle of similar developmental stage (right). Notice the reduced spacing between somatic and oocyte membranes in the *flam^Δ412-St2^* follicle. **(E)** Capsid accumulation (orange arrowheads pointing to several capsids) along the apical membrane of a somatic cell facing the nurse cells. **(F)** Capsid accumulations along the lateral membranes of two somatic cells, close to the apical side. For (C-F) Scale bar: 500 nm.

**Supplementary Figure 10:**
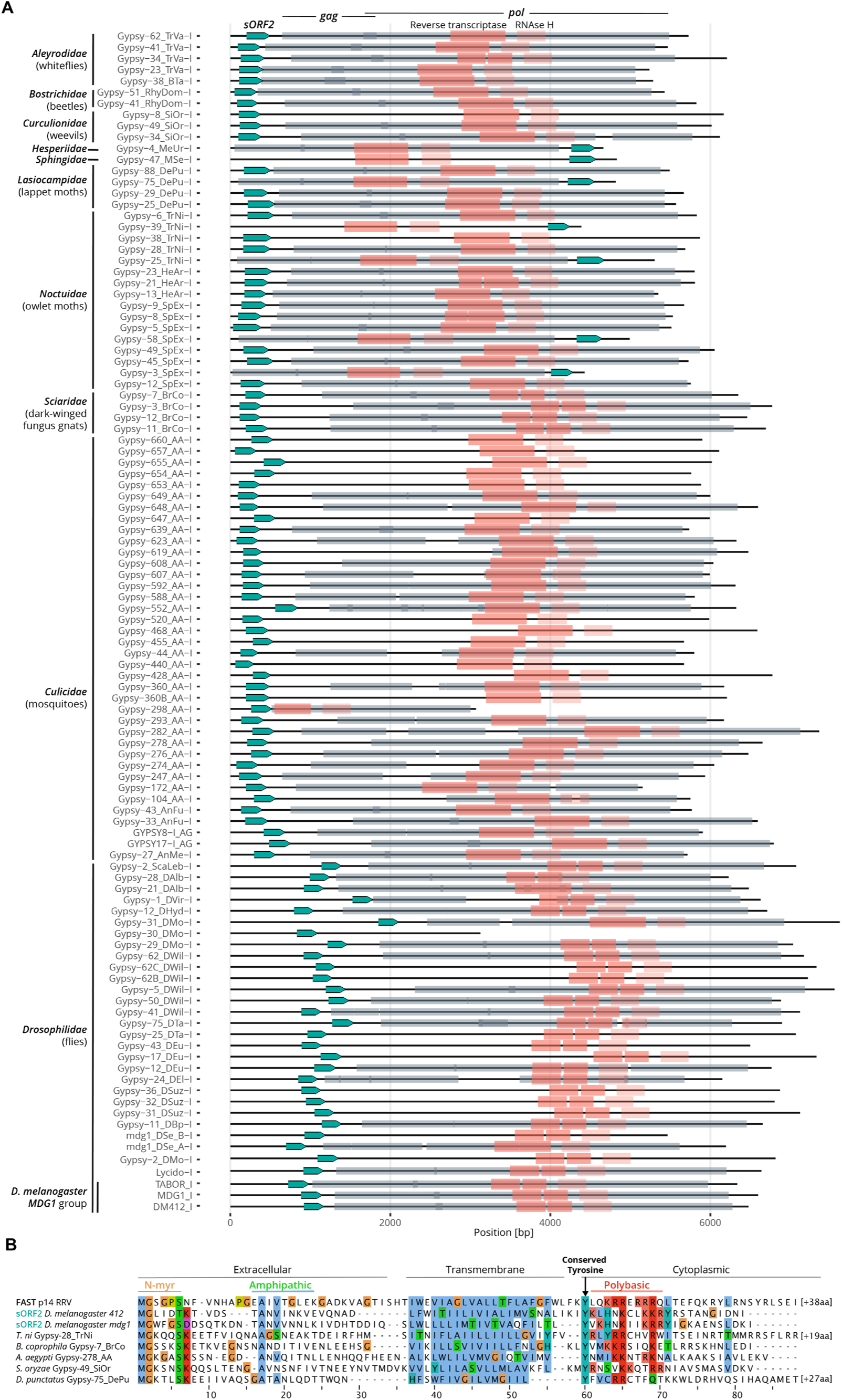
*sORF2/FAST*-like genes are encoded within *Metaviridae* in insect genomes. **(A)** Visual representation of internal consensus sequences from Repbase of all 105 identified *Gypsy* LTR retrotransposons with an *sORF2/FAST*-like gene (cyan arrow), across different insect families. Annotated ORFs from Repbase shown in grey boxes, likely corresponding to *gag* and *pol* ORFs. Identified regions with conserved domains reverse-transcriptase (dark pink) and RNAse H (light pink) support *pol* annotation. *LTR* regions are not shown. **(B)** Multiple sequence alignment of a known FAST protein p14, newly found sORF2 proteins from *D. melanogaster* and 5 representative sequences of sORF2/FAST-like proteins from insect *Metaviridae*. Notice conservation of all known structural features for FAST proteins and a newly identified Tyr residue just downstream of the TM.

**Supplementary Figure 11:**
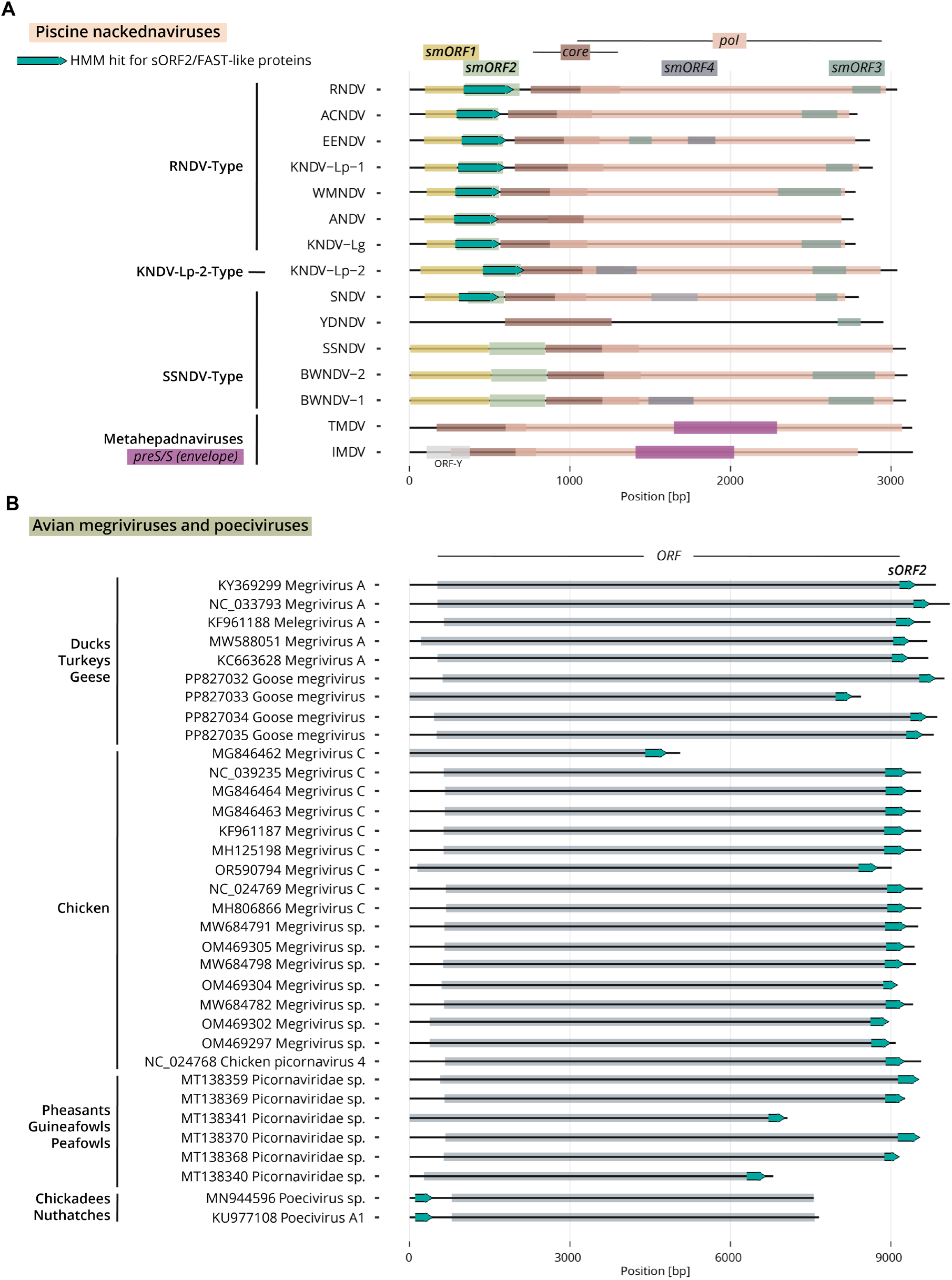
*sORF2/FAST*-like genes are encoded as separate ORFs within non-enveloped viral genomes. **(A)** Visual representation of assembled and annotated nackednavirus genomes according to (Lauber et al., 2017), with the putative smORF2 genes that were found by HMM-based search (cyan arrows). All other annotated ORFs are in colored boxes. SNDV and YDNDV have partial smORF2 sequences assembled. TMDV and IMDV are shown for comparison, with the annotated *preS/S* gene in violet **(B)** Visual representation of assembled and annotated megriviruses and poeciviruses from NCBI viral genomes (1,500-20,000 nt), with the putative *sORF2/FAST-like* ORF2 genes that were found by HMM-based search (cyan arrows). The hosts from which the viruses were sampled are noted on the left.

## Other supplementary materials for this manuscript

**Supplementary Table S1: *sORF2/FAST*-like sequences identified in this study.** The different tabs list sORF2/FAST-like sequences found in: (A) *Gypsy* LTR retrotransposon of insects. (B) *D. melanogaster*. (C) *A. aegyptii*. (D) *S. oryzae*. (E) Nackednaviruses. (F) Picornaviruses.

**Supplementary Table S2: Fly genotypes, smFISH probes and siRNA sequences used in this study.**

**Supplementary Movie S1: Electron tomography of an invasive protrusion in a *flam^Δ412-St2^* follicle.** The movie shows consecutive Z-sections from a tilt-series tomogram and a 3D rendering of an invasive protrusion containing retrotransposon capsids (pink), the somatic follicle cell membrane (green) and oocyte membrane (beige). Notice that the somatic and oocyte membranes are in extremely proximity, specifically at the site of the protrusion.

**Data S1:** High resolution spatial localization of capsid accumulations and invasive protrusions at the oocyte-soma interface in *flam^Δ412-St2^* follicle (same image as fig. S9, left)

**Data S2:** High resolution oocyte-soma interface in control follicle (same image as fig. S9, right)

