## Supplemental Data 1 for "Direct cell-to-cell transmission of retrotransposons"

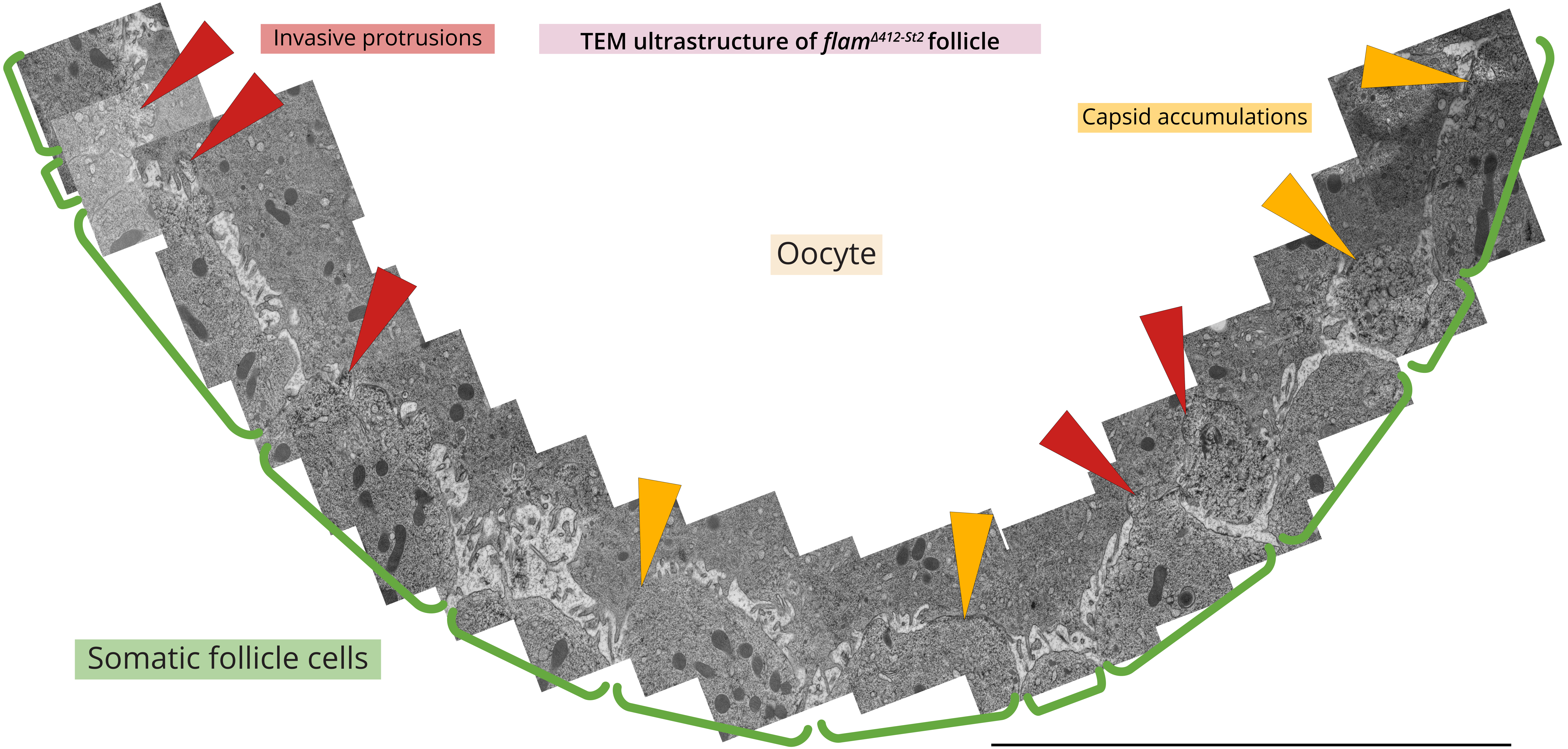


Data S1 (Voichek et al.): High resolution spatial localization of capsid accumulations and invasive protrusions at the oocyte-soma interface in *flam*$^{Δ412\text{-}St2}$ follicle (same image as fig. S9, left)
