## Supplementary material for "Direct cell-to-cell transmission of retrotransposons": Data S2: High resolution oocyte-soma interface in control follicle

### TEM ultrastructure of control follicle

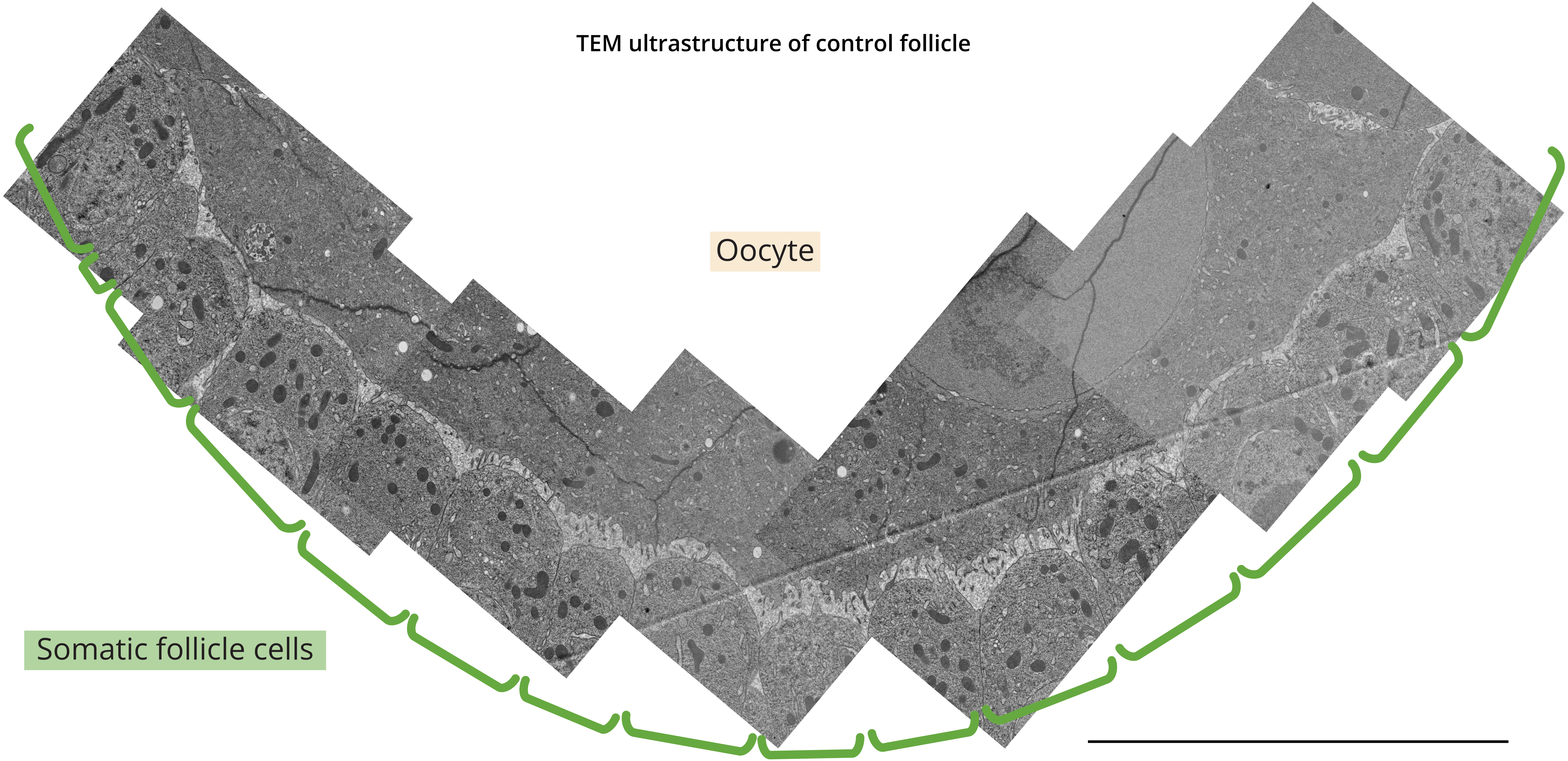


Data S2 (Voichek et al.): High resolution oocyte-soma interface in control follicle (same image as fig. S9, right)
